# FeatureMSEA: Metabolic Feature-based Metabolite Set Enrichment Analysis

**DOI:** 10.64898/2026.06.15.732313

**Authors:** Yijiang Liu, Yuting Wang, Tao Huan, Xiaotao Shen

**Author notes:** Corresponding author: Xiaotao Shen.

## Abstract

Liquid chromatography-mass spectrometry (LC-MS) untargeted metabolomics detects thousands of metabolic features, but converting these chemical signals into metabolite set-level biological knowledge remains challenging. This is because most features lack unambiguous metabolite identities. Conventional metabolite set enrichment analysis (MSEA) generally requires identified metabolites and metabolite-level ranked inputs, leaving much of the untargeted feature space unused. Here, we present FeatureMSEA, a feature rank-based framework for metabolite set enrichment directly from metabolic features with ambiguous annotations. FeatureMSEA integrates multi-evidence feature-to-metabolite annotation, feature rank-based enrichment scoring, permutation-based inference, and iterative leading-edge-guided annotation refinement, with an optional LLM-assisted module for post-enrichment interpretation. In null comparisons of randomly split healthy samples, FeatureMSEA detected no significant metabolite sets, whereas metabolite-set spike-in simulations showed recovery of implanted signals. In a cerebrospinal fluid metabolomics study of Huntington’s disease, FeatureMSEA identified dysregulated metabolite sets related to amino acid metabolism, mitochondrial energy metabolism, and neuroactive signaling. MS/MS-based annotation analysis further showed that FeatureMSEA refinement reduced annotation ambiguity and prioritized chemically consistent candidate metabolites. In summary, FeatureMSEA provides a general framework for extracting metabolite set-level biological insights from LC-MS untargeted metabolomics in which confident metabolite identification remains incomplete.

## Introduction

LC-MS untargeted metabolomics has become a central technology for profiling metabolic alterations associated with complex biological phenotypes, including disease, environmental exposure, microbiome activity, aging, diet, and drug response^1–4^. However, the primary output of LC-MS untargeted metabolomics is a metabolic feature table, in which each feature is typically defined by mass-to-charge ratio (*m/z*), retention time (RT), and abundance across samples^5^. Transforming these chemical signals into biological knowledge remains a major challenge^6^. This gap between metabolic features and interpretable biological entities limits the downstream functional analysis of untargeted metabolomics data^7^.

Despite its limitations, metabolite set analysis remains one of the most widely used strategies for interpreting metabolomics data^8^. Traditional over-representation analysis identifies metabolite sets that are statistically enriched among a subset of significant metabolites. However, this strategy requires users to define a significance threshold, and the results can be sensitive to arbitrary cutoffs^8^. In transcriptomics, gene set enrichment analysis (GSEA) addressed this limitation by testing whether members of a predefined gene set are non-randomly distributed along a ranked gene list, thereby avoiding the need to first select a subset of significant molecules^9^. This rank-based idea has also been extended to metabolomics as metabolite set enrichment analysis (MSEA)^10^. However, conventional MSEA usually assumes that the input can be represented as a list of confidently identified metabolites^10^. This assumption is often violated in LC-MS untargeted metabolomics, where confident metabolite identification is available for only a fraction of the metabolic features^11–13^.

Several approaches have shown that functional information can still be extracted from metabolic feature space even when metabolite identification is incomplete^14–16^. For example, mummichog infers metabolite-set activity from significant metabolic features with putative annotations^14^, PIUMet^15^ uses network-based strategies to identify disease-relevant metabolic modules, and FFMA from TidyMass2^16^ enables module-level interpretation directly from metabolic features. These methods have been important for relaxing the requirement of complete metabolite identification. Nevertheless, a distinct gap remains for a general, rank-based, GSEA-like metabolite set enrichment framework that operates directly on metabolic features with ambiguous annotations, without requiring prior selection of significant features^8,9,17^. Compared with methods that rely on significant feature subsets, such a framework should avoid arbitrary feature-level significance cutoffs by using the full phenotype-ranked feature list^18^. Ideally, such a method should preserve multiple candidate annotations per feature, perform metabolite set enrichment without requiring definitive metabolite identification, identify the feature-metabolite relationships that drive enrichment, and use this information to prioritize biologically plausible annotations.

Here, we present FeatureMSEA (Metabolic Feature-based Metabolite Set Enrichment Analysis), a rank-based framework for performing MSEA directly from metabolic features with ambiguous annotations. FeatureMSEA is based on the premise that biologically plausible annotations are more likely to make recurrent contributions to the leading edges of significantly enriched metabolite sets^9,14^. In contrast, incorrect candidate annotations are expected to behave more randomly and to appear less consistently among leading-edge contributors. We evaluate FeatureMSEA using simulated datasets with spike-in metabolite sets. We further apply FeatureMSEA to a cerebrospinal fluid metabolomics case study of Huntington’s disease to demonstrate disease-relevant metabolite set-level interpretation. Finally, we assess annotation refinement using confident MS/MS-supported annotations as partial reference evidence.

By explicitly modeling many-to-many feature-metabolite relationships, FeatureMSEA provides a feature-centric extension of MSEA for LC-MS untargeted metabolomics and enables metabolite set-level interpretation under annotation uncertainty. It is implemented as an accessible computational tool and can be integrated with existing metabolomics workflows, including TidyMass-compatible pipelines^16,19^.

## Results

### Overview of FeatureMSEA

The FeatureMSEA framework consists of five major steps: (1) feature-to-metabolite annotation, (2) feature rank-based enrichment scoring, (3) permutation-based significance assessment, (4) iterative annotation refinement, and (5) LLM-assisted biological plausibility assessment (**Fig. 1a-c**). Together, these steps enable FeatureMSEA to identify phenotype-associated metabolite sets directly from metabolic features.

**Figure 1.**
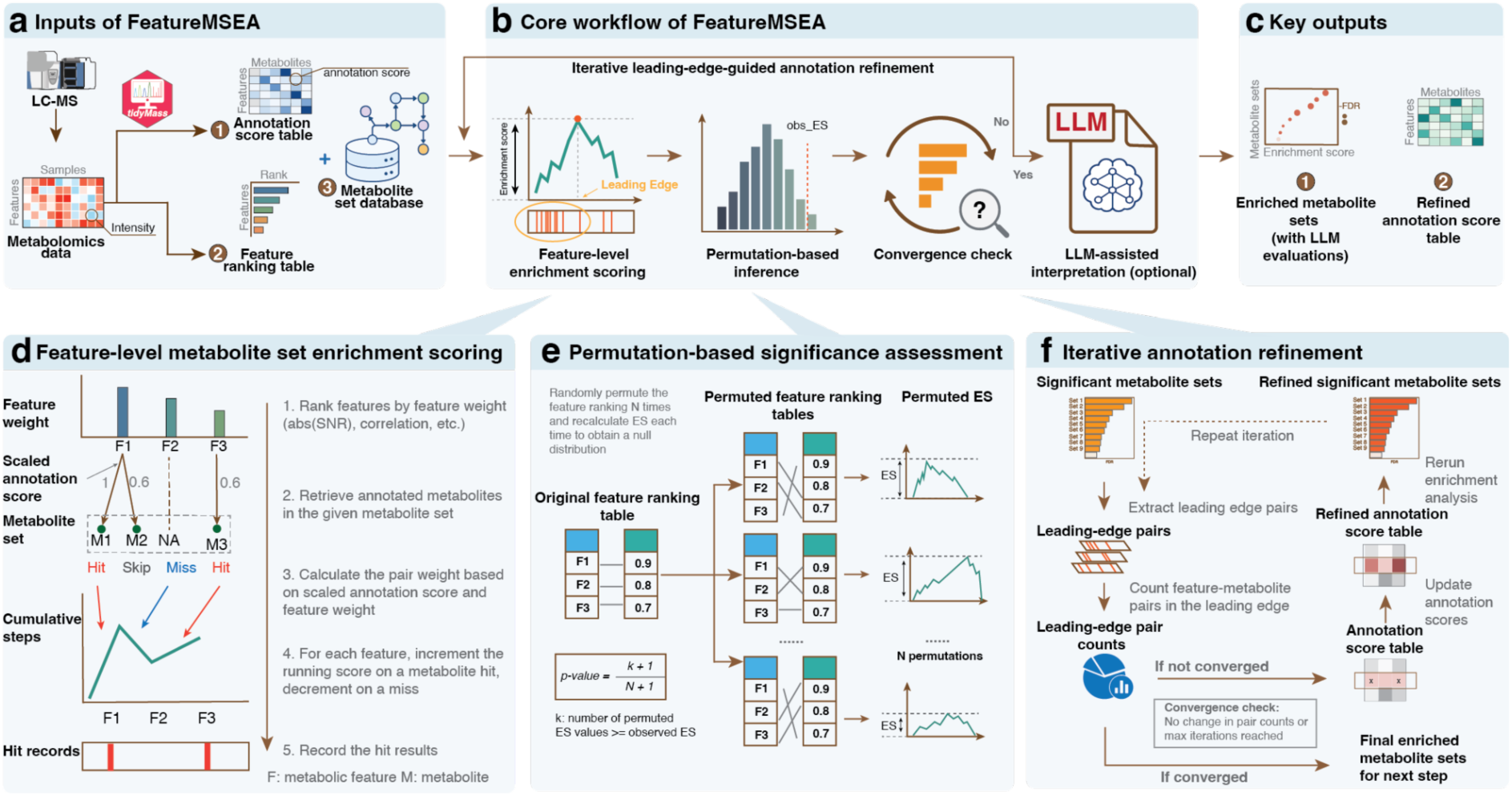
Workflow of FeatureMSEA. (a) Input data for FeatureMSEA. (b) Core FeatureMSEA workflow. (c) Main outputs of FeatureMSEA. (d) Feature-level enrichment scoring. Ranked features are matched to metabolites in a given metabolite set to compute a running enrichment score and record hit features. (e) Permutation-based inference. Feature ranking weights are permuted to generate null enrichment score distributions and estimate metabolite set-level significance. (f) Iterative annotation refinement. Leading-edge feature-metabolite pairs from significant metabolite sets are used to update annotation scores and rerun enrichment analysis until convergence.

The first step of FeatureMSEA is multi-evidence feature-to-metabolite mapping, which is adapted from the annotation strategy implemented in TidyMass2^16^ with minor modifications (**Fig. 1a**, **Methods**, and **Supplementary Note**). Briefly, starting from a feature table, features are annotated against a metabolite database using accurate mass matching, isotope patterns, adduct information, retention time, and, when available, MS/MS spectral similarity^20,21^. This step generates a feature-metabolite annotation table, in which each feature can be linked to one or more candidate metabolites with corresponding annotation scores. This annotation table serves as the basis for subsequent feature-level enrichment analysis (**Fig. 1a**).

The second step is feature-level metabolite set enrichment scoring (**Fig. 1d** and **Methods**). FeatureMSEA adapts the GSEA framework to the many-to-many relationship between features and metabolites^9^. Features are first ranked according to phenotype-associated weights (**Methods**). For each metabolite set, FeatureMSEA evaluates whether features linked to candidate metabolites in that set are non-randomly concentrated toward the top of the ranked feature list. Specifically, a running enrichment statistic is calculated across the ranked features. The statistic increases when a feature is associated with one or more metabolites in the tested metabolite set and decreases when no matched metabolite is present. The enrichment score (ES) is defined as the maximum deviation of this running statistic from zero (**Methods**). To account for annotation ambiguity, the contribution of each feature-metabolite pair is weighted by both the feature ranking weight and the scaled annotation score (**Methods**). In addition, FeatureMSEA applies a redundancy penalty to reduce over-representation by metabolites that are annotated to a large number of features. The final per-feature hit contribution is calculated from the penalized annotation score and the feature ranking weight and is normalized by the total hit contribution across all metabolite-set-associated features (**Fig. 1d**).

The third step is permutation-based significance assessment (**Fig. 1e** and **Methods**). To evaluate whether the observed enrichment score for each metabolite set is greater than expected by chance, FeatureMSEA constructs an empirical null distribution using feature-weight permutation. In each permutation, feature ranking weights are randomly reassigned across features, while the feature-metabolite annotation table is kept unchanged. The feature-level enrichment scoring procedure is then repeated for each permuted ranking table. By default, this permutation procedure is repeated 1,000 times for each metabolite set, generating a null distribution of enrichment scores. Empirical *P*-values are calculated by comparing the observed ES with the corresponding permuted ES distribution, and *P*-values are corrected for multiple testing using the Benjamini-Hochberg (BH) procedure^22^. Metabolite sets with an FDR below the user-defined threshold (default FDR < 0.05) are considered significantly enriched.

The fourth step is iterative leading-edge-guided annotation refinement, which uses metabolite set-level biological context to improve feature annotation prioritization (**Fig. 1f** and **Methods**). A key challenge in feature-based enrichment analysis is that feature-to-metabolite assignments are inherently uncertain, and different candidate annotations for the same feature can imply different metabolite-set memberships. FeatureMSEA addresses this challenge by identifying the leading-edge feature-metabolite pairs that drive significant metabolite set enrichment. These leading-edge pairs represent the feature-metabolite assignments that contribute most strongly to the observed enrichment signal. At each iteration, FeatureMSEA summarizes the recurrence and contribution of leading-edge feature-metabolite pairs across significant metabolite sets and uses this information to update the annotation weights. Candidate annotations that repeatedly contribute to the leading edges of significant metabolite sets are progressively reinforced, whereas annotations with little or no leading-edge support receive lower support in subsequent rounds. The updated annotation matrix is then used as input for another round of feature-level enrichment scoring and permutation-based significance assessment. Iteration continues until the set of significant feature-metabolite associations converges between consecutive rounds or until the maximum number of iterations is reached (**Methods**). This iterative refinement produces both refined metabolite-set enrichment results and a refined annotation score table, while preserving all candidate annotations rather than forcing each feature to a single metabolite identity (**Fig. 1f**).

The final step is LLM-assisted biological plausibility assessment, which provides an optional post-enrichment interpretation module (**Fig. 2** and **Methods**). This module does not determine statistical significance. Instead, it evaluates the biological plausibility of enriched metabolite sets by integrating enrichment results and relevant biological context. The detailed design and evaluation of this module are described in the next section (**Fig. 2**).

**Figure 2.**
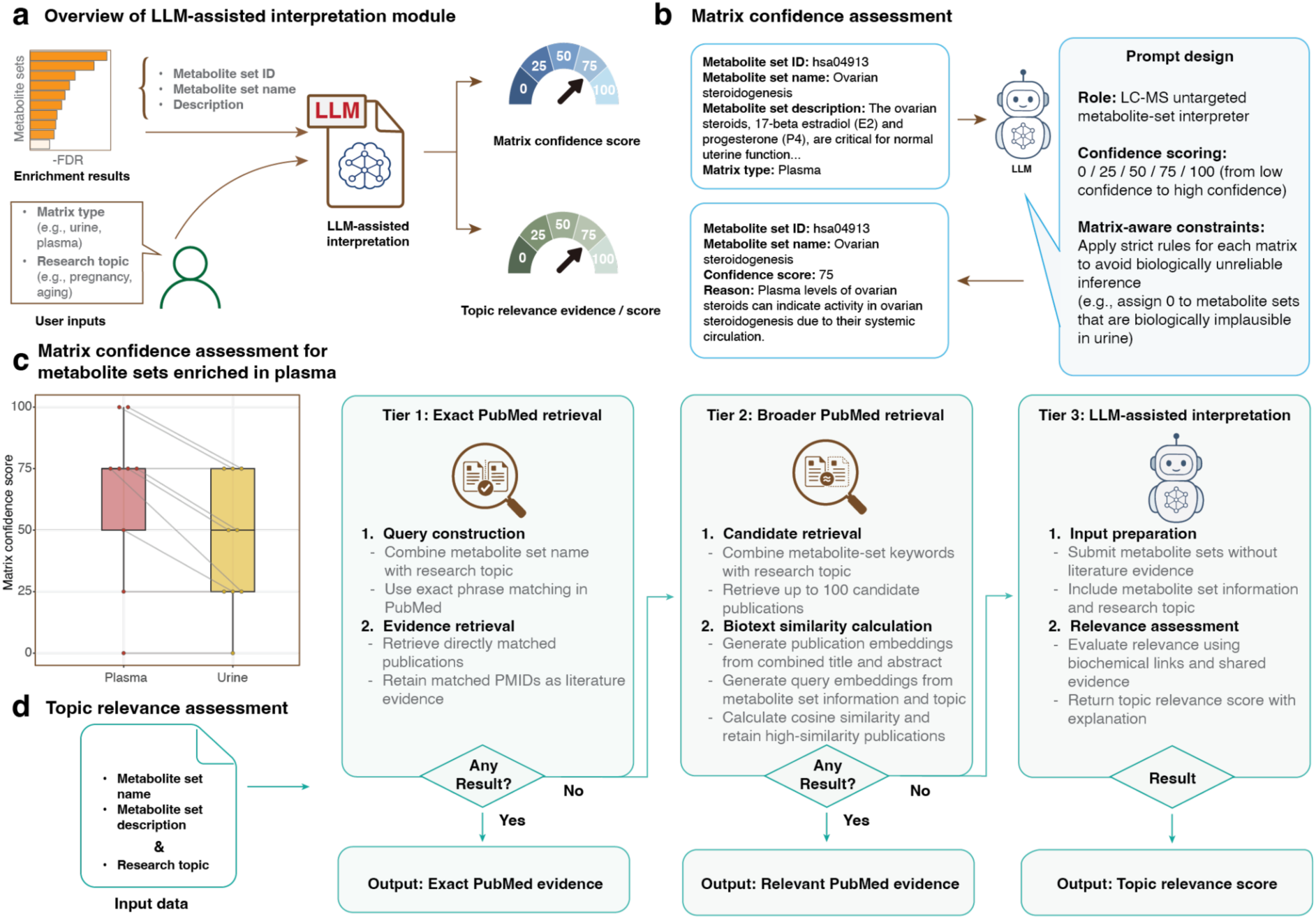
LLM-assisted biological plausibility assessment of enriched metabolite sets. (a) Overview of the LLM-assisted interpretation module. (b) Matrix confidence assessment. Metabolite set-level information is evaluated using a structured prompt with predefined scoring rules and matrix-aware constraints. (c) Evaluation of matrix confidence assessment using plasma-enriched metabolite sets. (d) Workflow of topic relevance assessment.

### LLM-assisted biological plausibility assessment of enriched metabolite sets

Although FeatureMSEA identifies statistically enriched metabolite sets from ranked metabolic features, statistical enrichment alone does not necessarily indicate that a metabolite set is biologically plausible or relevant within a specific experimental context^8^. This issue is particularly important in metabolomics, where metabolite set enrichment can sometimes produce statistically significant but biologically weak or poorly interpretable results^23^. To address this limitation, FeatureMSEA incorporates an LLM-assisted biological plausibility assessment module that evaluates enriched metabolite sets (**Fig. 2a** and **Methods**). This module functions as a post-enrichment interpretation layer and provides structured evidence to help users prioritize metabolite sets for downstream biological interpretation and experimental validation.

The LLM-assisted assessment module consists of two complementary components: (1) matrix confidence assessment and (2) topic relevance assessment (**Fig. 2a** and **Methods**). For matrix confidence assessment, FeatureMSEA provides the LLM with structured metabolite-set information together with the biological matrix type of the study (**Fig. 2b**). We designed a structured prompt consisting of three components: an overall task description, explicit scoring criteria, and strict output constraints^24,25^ (**Supplementary Note**). The LLM is instructed to evaluate whether the metabolite set represents a coherent and biologically plausible biochemical entity within the specified biological matrix. The output includes a matrix confidence score together with a concise explanation supporting the assigned score. This score is intended to estimate whether the enriched metabolite set is biologically reasonable within the given sample matrix.

To evaluate this module, we performed FeatureMSEA analysis on a plasma metabolomics dataset^26^ and assessed all enriched metabolite sets using the matrix confidence assessment workflow (**Methods**). First, the correct matrix information (plasma) was provided to the model, and a matrix confidence score was generated for each metabolite set. We then repeated the analysis using an intentionally incorrect matrix label (urine). As expected, most enriched metabolite sets received substantially higher confidence scores when the correct matrix information was provided compared with the incorrect matrix assignment (**Fig. 2c** and **Supplementary Table 1-2**). These results suggest that the matrix confidence assessment can capture matrix-specific plausibility of enriched metabolite sets.

The second component, topic relevance assessment, evaluates whether an enriched metabolite set is biologically relevant to a user-defined research topic. FeatureMSEA implements a three-tier evidence retrieval and evaluation strategy (**Fig. 2d** and **Methods**). In the first tier, FeatureMSEA performs an exact PubMed retrieval using the metabolite set name together with the user-defined research topic. This tier prioritizes directly published evidence linking the metabolite set to the biological topic.

If the exact search does not identify relevant literature, FeatureMSEA proceeds to a second tier involving broader PubMed retrieval (**Methods**). In this stage, individual terms are extracted from metabolite set names following preprocessing, then combined with the user-defined research topic to query up to 100 candidate publications. The titles and abstracts of all candidate publications are embedded and compared with a query vector constructed from the metabolite set and the user-defined research topic using cosine similarity^27,28^. Literature records with high similarity are retained as topic-relevant evidence.

If neither exact nor broad PubMed retrieval identifies relevant evidence, FeatureMSEA applies a third-tier fallback strategy based on LLM-assisted topic relevance scoring. In this mode, the LLM receives the research topic together with the metabolite set information and is instructed to evaluate the biological relevance of the metabolite set across four predefined dimensions: (1) known biochemical relationships, (2) upstream or downstream regulation, (3) shared metabolites or cofactors, and (4) previously proposed mechanistic hypotheses. The LLM then returns a structured topic relevance score together with a concise explanation. This fallback tier is only used when retrieval-based evidence is unavailable, thereby minimizing reliance on unconstrained generative inference and prioritizing direct literature-supported evidence whenever possible^29,30^.

### Metabolite-set spike-in simulations demonstrate recovery of implanted enrichment signals

To evaluate whether FeatureMSEA can recover metabolite sets with known dysregulated signals, we designed a metabolite-set spike-in simulation experiment using real LC-MS plasma metabolomics data (**Fig. 3a** and **Methods**). This simulation experiment was used to provide both a negative-control setting, in which no true biological difference was expected, and a positive-control setting, in which a predefined metabolite set was perturbed in silico and treated as the positive-control signal.

**Figure 3.**
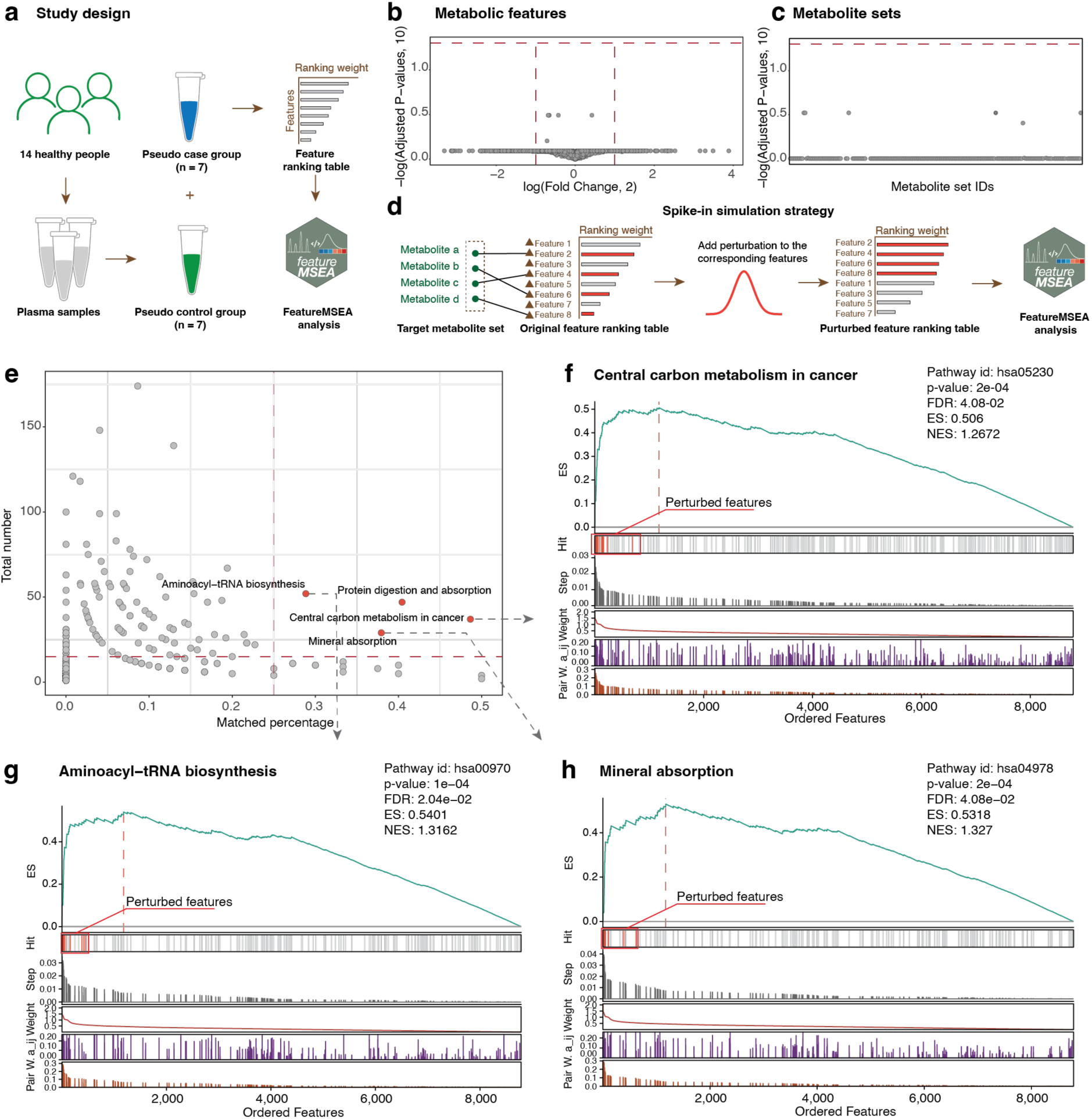
Metabolite-set spike-in simulations demonstrate recovery of implanted enrichment signals by FeatureMSEA. (a) Study design of the metabolite set spike-in simulation. (b) Differential feature analysis in the null comparison. (c) FeatureMSEA analysis in the null comparison. No metabolite sets were significantly enriched. (d) Strategy for metabolite set spike-in simulation. (e) Selection of target metabolite sets for spike-in simulation. (f-h) Enrichment plots for representative spike-in simulations. ES, enrichment score; NES, normalized enrichment score; FDR, false discovery rate.

We first selected plasma metabolomics data from 14 healthy individuals^31^ and randomly divided them into two pseudo groups, with 7 individuals assigned to each group (**Fig. 3a**). Because all samples were obtained from healthy individuals and group labels were randomly assigned, no systematic metabolic difference was expected between the two groups. As expected, differential feature analysis identified no significantly altered metabolic features (FDR < 0.05; **Fig. 3b**). We then applied FeatureMSEA to the same feature ranking table and found no significantly enriched metabolite sets at FDR < 0.05 (**Fig. 3c**). These results indicate that FeatureMSEA does not produce spurious metabolite set enrichment in a null comparison and support its specificity under a negative-control setting.

We next generated positive-control datasets by implanting known metabolite set-level signals into the same null background (**Methods**). Our rationale was that dysregulation of a metabolite set can be simulated by increasing the ranking weights of features corresponding to metabolites in that set^9,17,32^. For each target metabolite set, we first identified metabolic features annotated to metabolites^16,19^ in the selected set and then added normally distributed perturbation signals to their feature ranking weights, thereby creating a perturbed feature ranking table while keeping the original feature annotation structure unchanged (**Fig. 3d**). FeatureMSEA was then applied to the perturbed ranking table to test whether the implanted metabolite set could be recovered.

To select suitable target metabolite sets for spike-in simulation, we screened metabolite sets according to their coverage in the LC-MS metabolomics dataset (**Methods**). For each metabolite set, we calculated both the total number of metabolites in the set and the proportion of metabolites that could be mapped to features with MS/MS-based annotations. Metabolite sets with at least 15 metabolites and a mapped percentage of at least 25% were considered eligible for perturbation (**Fig. 3e**). This filtering ensured that each selected metabolite set had sufficient representation in the measured feature space to support a meaningful spike-in simulation.

Using these criteria, we selected four metabolite sets as positive-control targets, including central carbon metabolism in cancer, mineral absorption, aminoacyl-tRNA biosynthesis, and protein digestion and absorption (**Fig. 3e**). For each target set, perturbation signals were introduced separately into the corresponding features, and FeatureMSEA was run independently on each perturbed dataset. In three simulations, FeatureMSEA recovered the implanted metabolite set as the only significant enrichment result, without producing additional significant metabolite sets (**Fig. 3f-h** and **Supplementary Figure 1a-c**). In the remaining simulation, protein digestion and absorption was also significantly identified as the top-ranked enriched metabolite set. However, two additional metabolite sets were also detected as significant (**Supplementary Figure 1d and 2a-c**). Further inspection showed that these additional metabolite sets substantially overlapped with the implanted target set (**Supplementary Figure 2d-f**), indicating that they likely reflected shared metabolite membership rather than unrelated false-positive enrichment.

Together, these metabolite-set spike-in simulations show that FeatureMSEA behaves as expected in both negative-control and positive-control settings. These results support the ability of FeatureMSEA to identify dysregulated metabolite sets from ambiguously annotated metabolic features while maintaining low false-positive enrichment in null comparisons.

### FeatureMSEA reveals disease-relevant metabolic dysregulation in Huntington’s disease

To evaluate the ability of FeatureMSEA to identify disease-relevant metabolite set alterations in a real biological context, we applied it to an LC-MS untargeted metabolomics dataset of cerebrospinal fluid (CSF) from Huntington’s disease (HD) patients^33^. The dataset included CSF samples from 13 premanifest HD individuals and 13 manifest HD patients, providing a clinically relevant comparison for detecting metabolic alterations associated with disease progression (**Fig. 4a** and **Methods**). After data processing, a total of 6,331 metabolic features were detected and used for downstream analysis.

**Figure 4.**
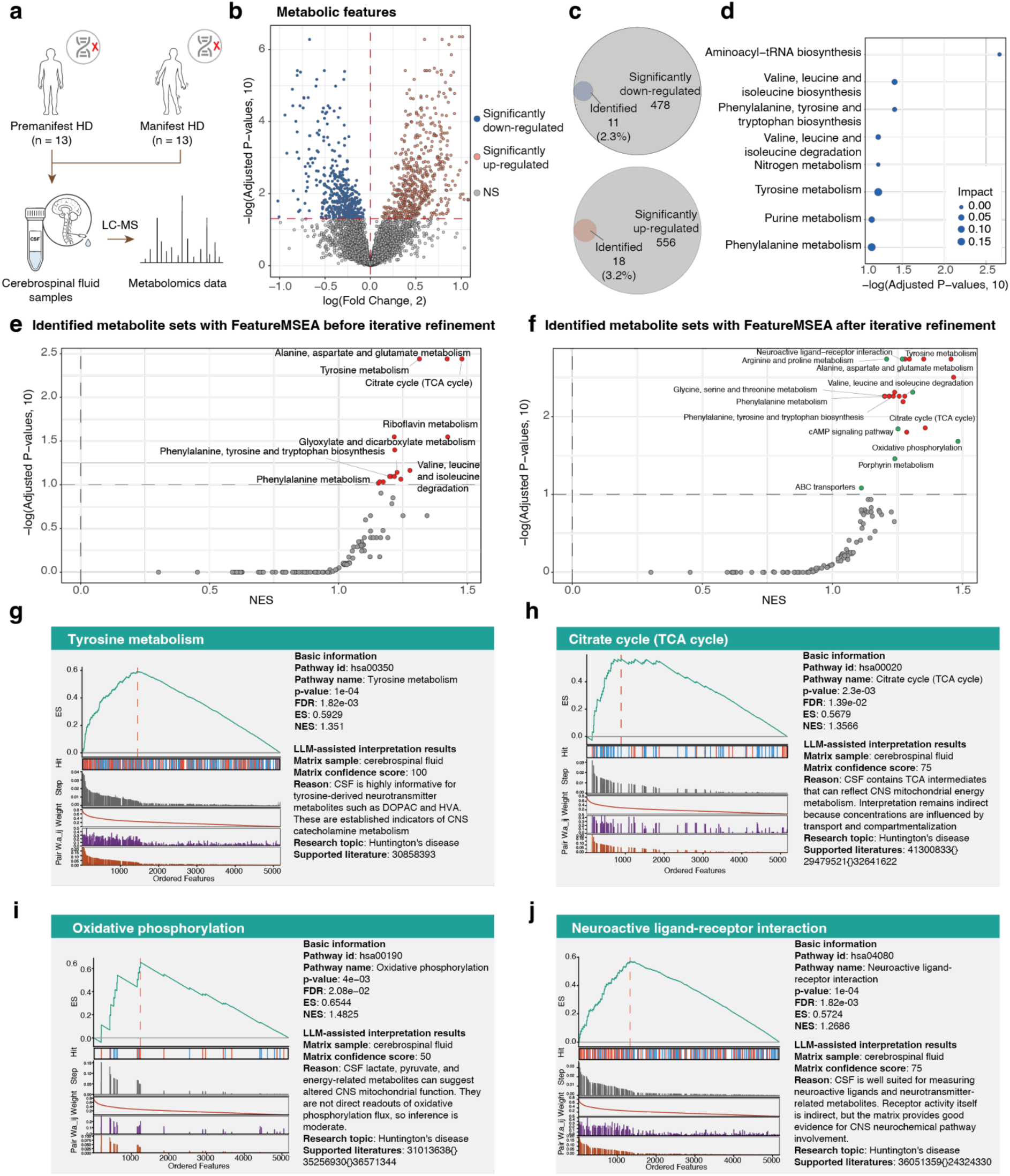
FeatureMSEA reveals disease-relevant metabolic dysregulation in HD. (a) Study design of the HD case study. (b) Volcano plot of metabolic feature alterations between the two groups. (c) Proportion of significantly altered features with metabolite-level annotations (MS/MS-based annotation) among upregulated and downregulated features. (d) Conventional metabolite-set analysis based on manually selected significant metabolites from the original study. (e) FeatureMSEA results before iterative annotation refinement. (f) FeatureMSEA results after iterative leading-edge-guided annotation refinement. (g-j) Representative enriched metabolite sets identified by FeatureMSEA. ES, enrichment score; NES, normalized enrichment score; FDR, false discovery rate; CSF, cerebrospinal fluid.

Differential feature analysis identified 556 significantly upregulated features and 478 significantly downregulated features in manifest HD compared with premanifest HD (**Fig. 4b**). However, only 3.2% of upregulated features and 2.3% of downregulated features had MS/MS-based metabolite annotations^16,19^, respectively (**Fig. 4c**).

In the original study, metabolite set analysis was performed using manually selected significant metabolites based on a predefined statistical threshold^33^. Such threshold-based analysis can be sensitive to the choice of significance cutoff, as illustrated in **Supplementary Figure 3**. Using this conventional enrichment strategy, 8 metabolite sets were reported as significantly enriched at FDR < 0.1 (**Fig. 4d**). We reasoned that FeatureMSEA could provide a complementary and more comprehensive analysis by using the full ranked feature list together with feature-metabolite annotation uncertainty.

We first applied FeatureMSEA without iterative annotation refinement. Using the complete phenotype-ranked feature list, FeatureMSEA identified 15 significantly enriched metabolite sets at FDR < 0.1 (**Fig. 4e** and **Supplementary Table 3**). These included metabolite sets related to amino acid metabolism, energy metabolism, and cofactor metabolism, several of which are consistent with known metabolic alterations in neurodegenerative disease^34,35^ (**Fig. 4e**). We then applied the full FeatureMSEA workflow with iterative leading-edge-guided annotation refinement. After refinement, FeatureMSEA identified 23 significantly enriched metabolite sets, including all 15 metabolite sets detected before refinement (**Fig. 4f** and **Supplementary Table 4**). This expansion of significant metabolite sets suggests that iterative refinement can strengthen metabolite set-supported feature-metabolite assignments and improve the detection of coherent metabolic dysregulation.

Among the enriched metabolite sets, tyrosine metabolism was one of the most prominent metabolite sets identified by FeatureMSEA (**Fig. 4g**). This result is consistent with the original report^33^ and with the known involvement of catecholamine-related metabolism in HD^34^. Tyrosine is a precursor of L-DOPA and dopamine, and dysregulation of this metabolite set may reflect altered dopaminergic neurotransmitter synthesis, turnover, or regulation^36,37^. Because dopaminergic dysfunction is closely linked to the motor and cognitive manifestations of HD, enrichment in tyrosine metabolism provides a biologically interpretable signal linking CSF metabolic changes to disease-related neuronal dysfunction^38,39^.

FeatureMSEA also identified metabolite sets related to mitochondrial energy metabolism, including the citrate cycle (TCA cycle) and oxidative phosphorylation (**Fig. 4h-i**). Dysregulation of the TCA cycle may indicate altered mitochondrial bioenergetic activity during HD progression, as this metabolite set supplies reducing equivalents required for ATP production through the electron transport chain^40,41^. The enrichment of oxidative phosphorylation further supports the involvement of mitochondrial dysfunction in manifest HD. Impaired oxidative phosphorylation can reduce ATP production and increase oxidative stress, both of which are consistent with neuronal vulnerability and disease progression in HD^42,43^. Together, these results suggest that FeatureMSEA captures coordinated metabolic alterations related to energy metabolism in the HD metabolome.

FeatureMSEA also identified neuroactive ligand-receptor interaction (**Fig. 4j**). Although not a canonical metabolic pathway, this metabolite set captures neuroactive small molecules and neurotransmitter-related signals^44–46^. We therefore interpreted this result as a neuroactive small-molecule signal rather than as direct evidence for receptor-level changes. This finding is consistent with the complex neurological phenotype of HD, in which motor, cognitive, and psychiatric symptoms may arise from widespread alterations in neuronal communication and synaptic regulation^47,48^.

The LLM-assisted biological plausibility assessment provided additional context for interpreting the enriched metabolite sets. For example, tyrosine metabolism received a high matrix confidence score in CSF and was supported by literature linking CSF catecholamine-related metabolites to HD-related neurochemical changes (**Fig. 4g**). The TCA cycle and oxidative phosphorylation were also prioritized as biologically plausible in the CSF context, consistent with mitochondrial energy dysfunction in HD^33,49^ (**Fig. 4h-i**).

Overall, this case study demonstrates that FeatureMSEA can identify biologically meaningful metabolite set dysregulation. FeatureMSEA recovered known HD-related metabolite sets and revealed additional metabolite sets related to amino acid metabolism, mitochondrial function, and neuroactive signaling. These findings support the utility of FeatureMSEA for metabolite set-level interpretation of disease-associated metabolic alterations in untargeted metabolomics datasets.

### MS/MS validation supports the reliability of FeatureMSEA-refined annotations

A key output of FeatureMSEA is the refined annotation score table, which prioritizes candidate feature-metabolite assignments using leading-edge information from significant metabolite sets. To evaluate whether this refinement improves annotation reliability, we assessed FeatureMSEA-refined annotations in the HD metabolomics dataset using MS/MS-based annotations as reference evidence (**Methods**).

We first extracted features contributing to the leading edges of significant metabolite sets and compared their candidate annotations before and after FeatureMSEA refinement. For each feature, candidate annotations were ranked according to either the original annotation score or the FeatureMSEA-refined annotation score. We then summarized the number of candidate annotations retained among the top-ranked positions. After FeatureMSEA refinement, the number of candidate annotations within the top 1, top 2, and top 3 ranked levels was markedly reduced compared with the original annotation table (**Fig. 5a**). This indicates that leading-edge-guided refinement reduces annotation ambiguity and prioritizes a smaller set of metabolite set-supported candidate metabolites, while preserving candidate-level uncertainty rather than forcing each feature to a single annotation.

**Figure 5.**
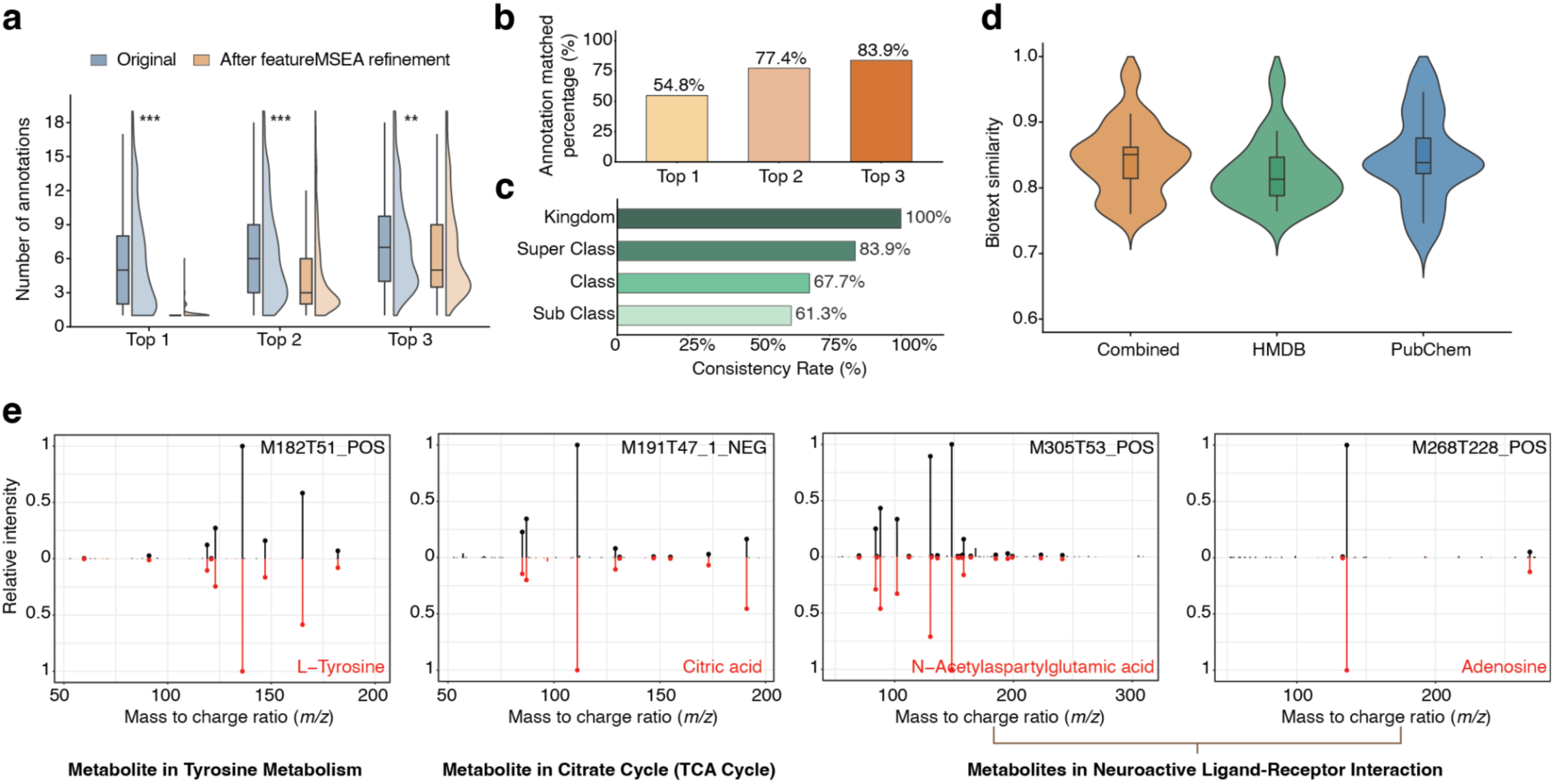
MS/MS validation supports the reliability of FeatureMSEA-refined annotations. (a) Comparison of annotation ambiguity before and after FeatureMSEA refinement. (b) Percentage of FeatureMSEA-refined annotations matched to MS/MS-based annotations at the top 1, top 2, and top 3 ranked levels. (c) Chemical taxonomy consistency between the top 1 FeatureMSEA-refined annotations and MS/MS-based annotations based on HMDB classification levels. (d) Biotext similarity between top 1 FeatureMSEA-refined annotations and the corresponding MS/MS-based annotations for non-exactly matched cases. (e) Representative MS/MS spectral matches supporting the top 1 FeatureMSEA-refined annotations.

We next evaluated whether the FeatureMSEA-prioritized annotations were supported by MS/MS evidence. Ranks were assigned using standard competition ranking, such that tied scores received the same rank and the next distinct score was ranked according to its position after accounting for ties. Using this criterion, 54.8% of the top 1 FeatureMSEA annotations matched MS/MS-based annotations, while the matched percentages increased to 77.4% and 83.9% when top 2 and top 3 annotations were considered, respectively (**Fig. 5b**).

Because multiple structurally related metabolites can produce similar MS/MS spectra or share close biological annotations^50–52^, we further assessed whether unmatched top 1 annotations were still chemically or biologically consistent with MS/MS-based annotations. For the top 1 FeatureMSEA annotations, we compared HMDB taxonomic classifications with the corresponding MS/MS-based annotations. The consistency rates were 100% at the kingdom level, 83.9% at the superclass level, 67.7% at the class level, and 61.3% at the subclass level (**Fig. 5c**). These results indicate that, even when exact annotation matches were not achieved, FeatureMSEA-prioritized annotations were frequently located within similar chemical categories to the MS/MS-based candidates.

To further evaluate semantic similarity between FeatureMSEA-prioritized annotations and MS/MS-based annotations, we retrieved metabolite descriptions from HMDB^53^ and PubChem^54^ and embedded these descriptions using an OpenAI embedding model (**Methods**). Cosine similarity was then calculated between each top 1 FeatureMSEA annotation and its corresponding MS/MS-based annotation for cases in which the top 1 FeatureMSEA annotation did not exactly match the MS/MS-based candidate. The resulting similarities were generally high across combined, HMDB-derived, and PubChem-derived descriptions (**Fig. 5d**), suggesting that many non-identical annotations prioritized by FeatureMSEA remain biologically or chemically related to the MS/MS-based annotations.

Finally, we inspected representative MS/MS spectral matches for the top 1 FeatureMSEA annotations that were also supported by MS/MS evidence. Four examples, L-tyrosine, citric acid, N-Acetylaspartylglutamic acid, and adenosine, showed consistent fragmentation patterns between the experimental MS/MS spectra and the reference MS/MS spectra of the corresponding compounds (**Fig. 5e**). Other annotations are provided in **Supplementary Figure 4**.

Together, these analyses support the reliability of the FeatureMSEA-refined annotation table. Leading-edge-guided refinement reduced the number of competing candidate annotations, increased the prioritization of MS/MS-based candidates, and retained chemical or semantic consistency even when the exact top 1 annotation did not match the MS/MS-based annotation. These results demonstrate that FeatureMSEA not only identifies dysregulated metabolite sets but also improves the interpretability of feature-level annotations by prioritizing metabolite set-supported and MS/MS-consistent candidate metabolites.

## Discussion

In this study, we developed FeatureMSEA, a metabolic feature-based framework that extends MSEA to LC-MS untargeted metabolomics with incomplete and ambiguous metabolite annotations. A central challenge in untargeted metabolomics is that the primary analytical output is a feature table rather than a confidently identified metabolite list. Conventional MSEA generally requires identified metabolites, which are often unavailable for the majority of metabolic features. FeatureMSEA addresses this gap by integrating multi-evidence feature-to-metabolite annotation, feature rank-based enrichment scoring, permutation-based significance assessment, iterative leading-edge-guided annotation refinement, and LLM-assisted biological plausibility assessment into a unified workflow. By retaining multiple candidate annotations for each feature and propagating them into a rank-based enrichment framework, FeatureMSEA enables metabolite set-level interpretation without forcing each feature to a single metabolite identity.

A key methodological innovation of FeatureMSEA is its iterative leading-edge-guided annotation refinement. Annotation uncertainty is intrinsic to LC-MS untargeted metabolomics. Rather than discarding ambiguous features or selecting one annotation before analysis, FeatureMSEA uses the enrichment structure itself to prioritize candidate feature-metabolite assignments. In this framework, leading-edge feature-metabolite pairs represent the assignments that contribute most strongly to significant metabolite set enrichment. This strategy prioritizes candidate annotations that are more consistent with the phenotype-associated metabolite set-level signal.

FeatureMSEA also incorporates an LLM-assisted biological plausibility assessment module to support post-enrichment interpretation. Statistical enrichment alone does not guarantee that a metabolite set is biologically meaningful in a given experimental context^55,56^. The LLM-assisted module addresses this challenge through matrix confidence assessment and topic relevance assessment. The LLM-assisted module does not determine enrichment significance, replace expert review, or validate biological mechanisms. Instead, it helps organize evidence, rank biological plausibility, and guide users toward enriched metabolite sets that are more likely to be interpretable in the context of the study.

Several limitations should be considered when applying FeatureMSEA. First, FeatureMSEA is identification-independent but not annotation-free. It requires candidate feature-metabolite mappings, and its performance depends on the quality, coverage, and accuracy of the annotation table. Second, leading-edge recurrence can be influenced by metabolite set overlap and hub metabolites that appear in many metabolite sets. Although FeatureMSEA incorporates redundancy penalties to reduce over-representation, more explicit correction for metabolite set degree or hub-metabolite effects may further improve annotation refinement. Third, FeatureMSEA currently uses magnitude-based ranking and is designed to identify dysregulated metabolite sets rather than directionally upregulated or downregulated metabolite sets. This design is biologically motivated because metabolites within the same metabolite set can change in opposite directions due to substrate-product relationships, branch-point regulation, or compensatory metabolic fluxes^23^. Nevertheless, some applications may benefit from optional signed enrichment analyses in future versions. Finally, the LLM-assisted assessment requires careful interpretation. LLM outputs depend on prompt design, model behavior, and available literature, and the resulting scores should be used to support, not replace, expert interpretation or experimental validation^24,57^. Future studies could benchmark LLM-assisted metabolite set interpretation against expert-curated relevance annotations.

In summary, FeatureMSEA provides a general and extensible framework for transforming ambiguously annotated metabolic features into interpretable metabolite set-level biological insights. FeatureMSEA bridges the gap between untargeted metabolomics feature tables and metabolite set-level interpretation. This framework should be broadly useful for disease metabolomics, exposure biology, drug response profiling, and other LC-MS untargeted metabolomics applications in which confident metabolite identification remains incomplete.

## Methods

### Feature-based metabolite set enrichment analysis

FeatureMSEA takes a metabolic feature table as the primary input and proceeds through five sequential steps: multi-evidence feature-to-metabolite annotation, feature rank-based enrichment scoring, permutation-based significance assessment, iterative leading-edge-guided annotation refinement, and LLM-assisted biological plausibility assessment. We provide an R package and a web interface at https://github.com/tidymass/featuremsea and https://featuremsea.jaspershenlab.com/.

#### Data preparation

The required input feature table includes variable_id (unique identifier), *m/z*, rt (retention time in seconds), condition (phenotype-associated ranking statistic such as absolute signal-to-noise ratio (SNR), absolute log2 (fold change) (log2FC), and absolute correlation coefficient), polarity mode (“positive” or “negative”), and mean_intensity (average feature intensity across samples). The analysis additionally requires an MS1 metabolite database and a metabolite set database. The MS1 metabolite database can be selected from KEGG^45^ or HMDB^53,58^, and the metabolite set database can be selected from five publicly available metabolite set databases: KEGG^45^, PathBank^59^, MetaCyc^60^, WikiPathways^61^, and Reactome^62^. To mitigate redundancy across these resources, we constructed an integrated metabolic pathway database (iMetPD) by consolidating entries from all five sources (**Supplementary Figure 5** and **Supplementary Note**). All curated public databases are freely accessible through the TidyMass website: https://www.tidymass.org/databases/.

#### Multi-evidence feature-to-metabolite annotation

The feature-to-metabolite annotation procedure in FeatureMSEA is adapted from the annotation strategy implemented in TidyMass2^16^. First, all metabolic features are matched to candidate metabolites based on accurate mass within a user-defined mass tolerance (default: 15 ppm). Second, metabolic features are grouped into a metabolite feature cluster (MFC) based on chemical identity and chromatographic behavior. Third, a confidence score is assigned to each MFC based on chemical evidence such as theoretical isotope patterns and characteristic adducts (**Supplementary Table 5** and **Supplementary Note**). Fourth, redundant annotations are removed based on confidence score to reduce false positive rates. The detailed information is provided in the **Supplementary Note**.

#### Feature rank-based enrichment scoring

Features are first ranked in descending order based on the absolute value of the phenotype-associated ranking statistic *x_i_*, such as SNR, log2FC, and correlation coefficient. For two-group comparisons, SNR is used as the recommended statistic^63^ and computed as:

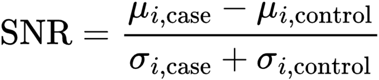

where *µ_i,case_* and *µ_i,control_* represent the mean log_2_(*x+1*) transformed intensities of feature *i* in the case and control groups, respectively, and *σ_i,case_* and *σ_i,control_* represent the corresponding within-group standard deviations. The ranking weight of feature i is then defined as:

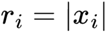

For each metabolite set, FeatureMSEA evaluates whether features matched to the metabolites concentrate at the top of the feature ranking table instead of forming a random distribution, following the general logic of GSEA^9^. For each tested metabolite set, a feature is considered a “Hit” if it is annotated to at least one metabolite in that set. When multiple candidate metabolites from the same set are assigned to the same feature, only the candidate with the highest annotation weight is used for the enrichment score calculation to avoid repeated contribution of a single feature. Other candidate annotations are retained in the output. If a feature cannot be annotated to any metabolite within the given metabolite set, it is designated as a “Miss”. The annotation weight *a_i,j_* between feature i and metabolite j is defined as:

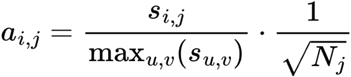

where *s_i,j_* represents the original enrichment annotation score between feature i and metabolite j, *max_u,v_*(*s_u,v_*) indicates the global maximum value across the entire annotation score matrix, and *N_j_* is the number of features annotated to metabolite j, which serves as a penalty coefficient to reduce over-representation by metabolites that are matched to a large number of features.

For each metabolite set M, a running enrichment score is computed by traversing the ranked feature list from top to bottom. At each position i, the enrichment score accumulates at a “Hit” position:

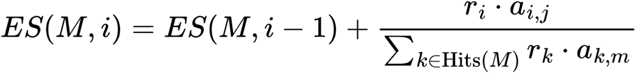

the enrichment score decreases at a “Miss” position:

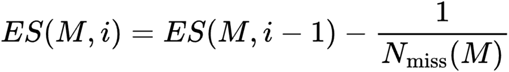

where *N_miss_* (*M*) is the number of features not annotated to any metabolite in metabolite set M. The enrichment score (ES) is defined as the maximum absolute deviation of the running statistic from zero:

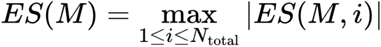

#### Permutation-based inference

To evaluate the statistical significance of the observed enrichment score for each metabolite set, FeatureMSEA constructs a null ES distribution by permuting feature ranking weights. In each permutation, the ranking weights are randomly reassigned across features while the feature-metabolite annotation table is unchanged. The enrichment score procedure is repeated for *N_prem_* times (default: 1,000) to obtain a null distribution of enrichment scores. The normalized enrichment score (NES) is then computed by scaling the observed ES by the mean of the null distribution. For each metabolite set, the *P-*value is defined as:

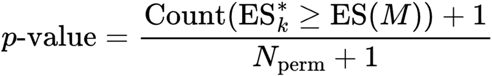

*P-values* are further adjusted for multiple testing across all tested metabolite sets using the BH procedure. Metabolite sets with the FDR below the user-defined threshold (default: FDR < 0.05) are considered significantly enriched.

#### Iterative leading-edge-guided annotation refinement

To refine feature-metabolite annotation scores using biologically supported leading-edge evidence, FeatureMSEA iteratively updates the annotation scores based on leading-edge information from significantly enriched metabolite sets. At each iteration, the leading-edge feature-metabolite pairs are retrieved from all significant metabolite sets. For each feature-metabolite pair (i, j), a recurrence count *c_i,j_* is calculated to quantify the number of significant metabolite sets in which the pair occurs within the leading edge. The bonus for each feature-metabolite pair (i, j) is then defined as:

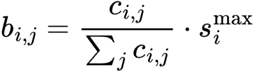

where 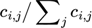 represents the feature-based occurrence proportion, and 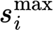 serves as the feature-specific scaling baseline to ensure dimensional consistency. The refined annotation score is then computed as:

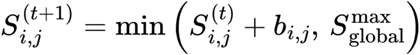

where 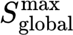 represents the maximum annotation score across all feature-metabolite pairs in the annotation score table, which serves as an upper bound to limit the impact of extreme score values on subsequent enrichment scoring. For feature–metabolite pairs that do not appear in the leading edge, no bonus is added, and the annotation score remains unchanged.

The refined annotation score table is then used as input for the next round of feature rank-based enrichment scoring and permutation-based inference. The iterative procedure is repeated until either the counts of feature-metabolite pairs in the leading edge remain unchanged between two consecutive iterations or the maximum number of iterations is reached (default: 3).

#### LLM-assisted biological plausibility assessment

To support post-enrichment biological interpretation, FeatureMSEA incorporates an optional LLM-assisted module to evaluate the biological plausibility of significant metabolite sets obtained from previous steps without affecting any statistical indicators. The module includes matrix confidence assessment and topic relevance assessment.

For matrix confidence assessment, basic information of all significant metabolite sets is submitted to the LLM in a single batch call. Each metabolite set is evaluated using a structured prompt that incorporates the metabolite set identifier, metabolite set name, and the biological matrix type of the study (**Supplementary Note**). A matrix confidence score on a discrete five-point scale (0, 25, 50, 75, or 100), accompanied by a brief explanatory rationale, is returned for the assessment. The underlying LLM backend is user-configurable, with support for OpenAI (default: GPT-5.5) and SiliconFlow (default: Qwen3-32B).

For topic relevance assessment, FeatureMSEA uses a three-tier strategy that prioritizes literature-based evidence over LLM inference. An exact PubMed retrieval is first performed using the user-defined research topic and the metabolite set name from the significant metabolite set. If no exact matches are found, candidate publications are retrieved using keywords from the metabolite set name combined with the research topic. Titles and abstracts of the top 100 relevant publications are embedded using a selectable embedding model provided by OpenAI (default: text-embedding-3-small) or SiliconFlow (default: Qwen3-Embedding-8B) and ranked by cosine similarity to a query vector constructed from the metabolite set name, description, and research topic. Publications with similarity scores above the threshold (default: 0.6) are retained. If no relevant literature evidence is identified, an LLM-assisted fallback is used to assign a topic relevance score on the same five-level scale based on known biochemical links between the metabolite set and the research topic, potential upstream or downstream regulation, shared metabolites or cofactors, and published hypotheses or emerging evidence (**Supplementary Note**).

### Matrix confidence assessment validation

To validate the reliability of the matrix confidence assessment module in LLM-assisted biological plausibility assessment, we applied FeatureMSEA to an LC-MS plasma metabolomics dataset from a public longitudinal pregnancy cohort^26^. The dataset comprised 784 weekly plasma samples collected from 30 healthy pregnant women spanning from gestational week 5 through the postpartum period. Raw LC-MS data were processed using TidyMass^19^, yielding 13,314 metabolic features. The correlation coefficient between each feature’s abundance and gestational age was adopted as the ranking statistic.

The resulting feature ranking table was used as the primary input for FeatureMSEA, which identified 9 significantly enriched metabolite sets from the KEGG human metabolite set database (1,000 permutations; FDR < 0.05). Matrix confidence assessment was then applied to all 9 metabolite sets using GPT-5.5 with either the correct matrix label (plasma, **Supplementary Table 1**) or an intentionally incorrect label (urine, **Supplementary Table 2**).

### Metabolite-set spike-in simulation

To evaluate FeatureMSEA performance under spike-in simulation, we analyzed a publicly available LC-MS plasma metabolomics dataset derived from 14 healthy individuals^31^. The 14 individuals were randomly divided into two pseudo-groups of 7 (simulated case and control). Raw data were processed using TidyMass^19^. Differential feature analysis was performed for each metabolic feature by calculating the log2FC between simulated case and control groups and conducting two-sample t-tests. The resulting *P*-values were adjusted for multiple testing using the Benjamini–Hochberg method, and features with an FDR < 0.05 were considered significant. FeatureMSEA was then applied to the feature table using the absolute SNR as the ranking statistic. The SNR was calculated between the simulated case and control groups, and enrichment significance was evaluated using 10,000 permutations with an FDR threshold of 0.05.

For positive-control simulations, features were first annotated based on MS/MS spectra using TidyMass^19^. For each metabolite set in the KEGG human pathway database^58,64^, we calculated the proportion of metabolites in the set that were matched by at least one MS/MS-annotated feature. Metabolite sets with a matched percentage of at least 25% and a total compound count of at least 15 were selected as spike-in targets. For each target set, perturbation signals sampled from N(1, 0.25^2^) were added to the ranking weights of features annotated to metabolites in the target set, generating a perturbed feature ranking table while keeping the metabolite annotation table unchanged. FeatureMSEA was then applied to each perturbed feature table independently using 10,000 permutations and an FDR threshold of 0.05 to test whether the implanted metabolite set could be recovered.

### Huntington’s disease case study

This study reanalyzed publicly available LC-MS metabolomics data from cerebrospinal fluid samples of a Huntington’s disease cohort; no new metabolomics data were generated^33^. LC-MS profiles from 13 premanifest and 13 manifest HD individuals were designated as the control and case groups, respectively. Raw data were processed using TidyMass^19^. Differential feature analysis was performed for each metabolic feature by calculating the log2FC between manifest and premanifest HD groups and conducting two-sample t-tests. The resulting *P*-values were adjusted for multiple testing using the BH procedure, and features with an FDR < 0.05 were considered significant.

All 6,331 detected metabolic features were included in the FeatureMSEA analysis without pre-filtering for statistical significance. Features were ranked by the absolute SNR calculated between the manifest and premanifest HD groups. FeatureMSEA was applied under two settings: a non-refined setting with a single enrichment run, and an iterative refinement setting with a maximum of three refinement iterations. Both analyses used 10,000 permutations, with an FDR threshold of < 0.1 for the identification of enriched metabolite sets.

LLM-assisted biological plausibility assessment was applied to evaluate the enriched metabolite sets. For matrix confidence assessment, GPT-5.5 was queried with “cerebrospinal fluid” as the sample source keyword. Topic relevance assessment was conducted using “Huntington’s disease” as the research topic, with semantic embeddings generated by the text-embedding-3-small model.

### MS/MS-based annotation validation

MS/MS spectra data from the Huntington’s disease cerebrospinal fluid metabolomics dataset were used to evaluate FeatureMSEA annotation results. MS/MS-based metabolite annotation was performed with TidyMass^19^. We extracted all 496 features that appeared in the leading edge of significant metabolite sets after iterative refinement of FeatureMSEA. Candidate metabolites in the feature-metabolite annotation score table were ranked according to their annotation scores. Candidate metabolites were ranked separately using the original annotation score table and the FeatureMSEA-refined annotation score table. The number of candidate annotations retained within the top 1, top 2, and top 3 ranked positions was then calculated for each feature. Standard competition ranking was used to handle tied scores.

Among the 496 features, 31 features were found to have MS/MS-based annotations. For each of them, we assessed whether the MS/MS-based annotation was recovered among the top 1, top 2, or top 3 FeatureMSEA-ranked candidates. The top-k matching rate was calculated as the proportion of MS/MS-based leading-edge-associated features whose reference annotation was recovered within each rank threshold. For the top 1 annotations, we further obtained their corresponding classification information based on the HMDB database^53^ and calculated the consistency rate, including kingdom, superclass, class, and subclass. For the top 1 FeatureMSEA annotations that did not exactly match the corresponding MS/MS-based annotations, HMDB^53^ and PubChem^54^ descriptions were retrieved and embedded for both metabolites. Cosine similarity between the resulting embeddings was calculated to quantify biotext similarity. For exactly matched top 1 annotations, MS/MS spectra were retrieved and further compared with reference MS/MS spectra of the corresponding standard compounds.

## Data Availability

The LC-MS metabolomics data analyzed in this study were obtained from previously published studies^26,31,33^. The metabolomics data used in the matrix confidence assessment have been deposited in the Metabolomics Workbench under Study ID <u>PR000918</u>. The metabolomics data used in the metabolite-set spike-in study have been deposited in the Metabolomics Workbench under Study ID <u>ST002796</u>. The metabolomics data used in the HD study have been deposited in the MetaboLights under Project ID <u>MTBLS749</u>.

## Code Availability

This study utilized R version 4.5.2 and its associated packages for software development, data processing, and statistical analysis. The complete source code for FeatureMSEA is freely available at https://github.com/tidymass/featuremsea. The FeatureMSEA Shiny application is available at https://featuremsea.jaspershenlab.com/, and its source code is freely available at https://github.com/tidymass/featuremsea_shiny. All the databases (metabolite set databases and MS1 metabolite databases) are available at https://www.tidymass.org/databases/. All code used for data processing, analysis, and visualization in this study can be accessed at https://github.com/tidymass/fmsea_manuscript.

## Author Contributions Statement

X.S. conceived the method and provided overall supervision. Y.L. and X.S. jointly developed the methods. Y.L. implemented FeatureMSEA and developed its Docker version. Y.L. prepared documentation and tutorials. Y.L. analyzed the case study data. Y.L. generated all figures. Y.L., Y.W., and X.S. wrote the manuscript. T.H. reviewed and contributed to the final manuscript.

## Acknowledgments

This work was supported by start-up funding (Nanyang Assistant Professorship) provided to Dr. Xiaotao Shen from the Lee Kong Chian School of Medicine (LKCMedicine) and the School of Chemistry, Chemical Engineering, and Biotechnology (CCEB) at Nanyang Technological University, Singapore (NTU). This work was also supported by a Tier 1 grant from the Ministry of Education (MOE), Singapore, to Xiaotao Shen (Grant number: #025402-00001). The authors acknowledge the use of ChatGPT (OpenAI) for assistance with language polishing and grammar correction during the preparation of this manuscript.

