## Supplementary information for "FeatureMSEA: Metabolic Feature-based Metabolite Set Enrichment Analysis"

### Supplementary Figures

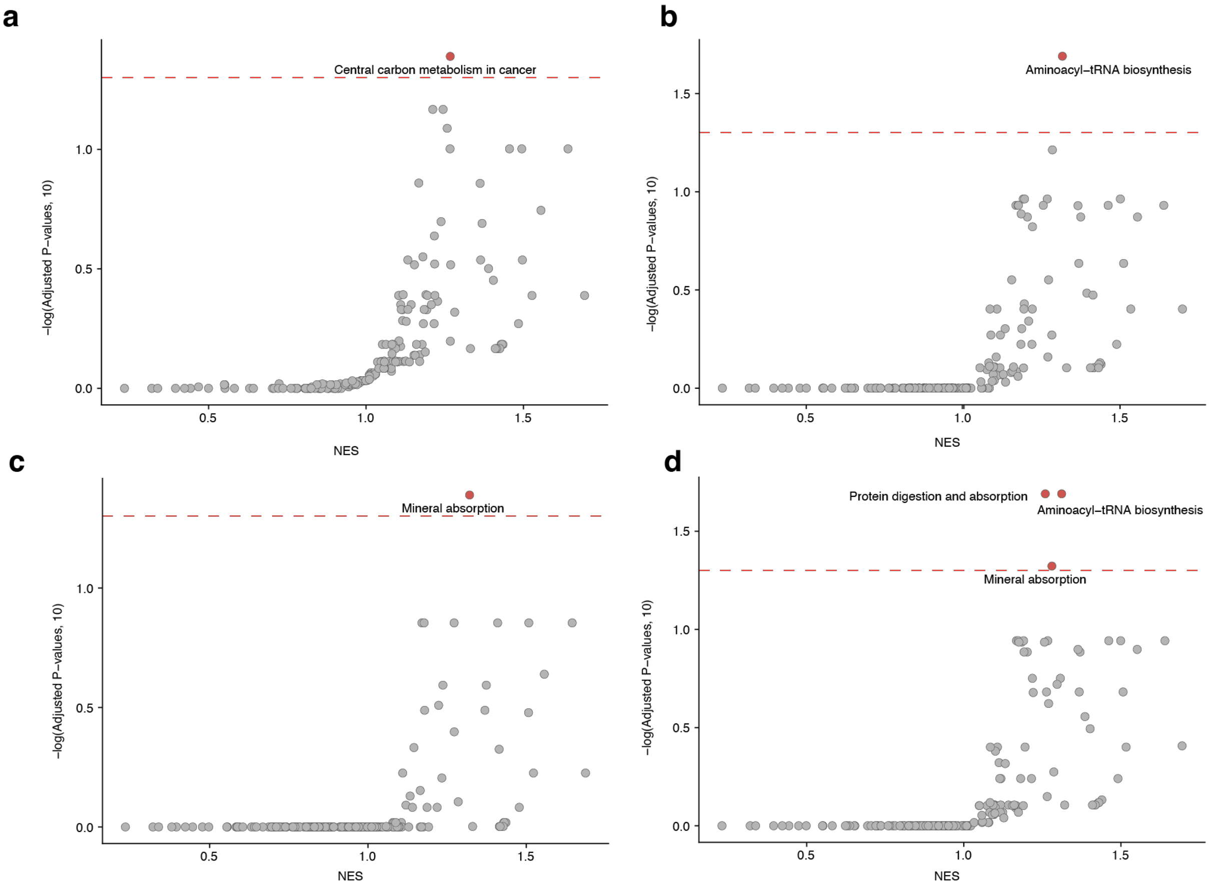


**Supplementary Figure 1 | Enrichment profiles of spike-in simulations across four positive-control metabolite sets. (a)** Central carbon metabolism in cancer. **(b)** Aminoacyl−tRNA biosynthesis. **(c)** Mineral absorption. d, Protein digestion and absorption.

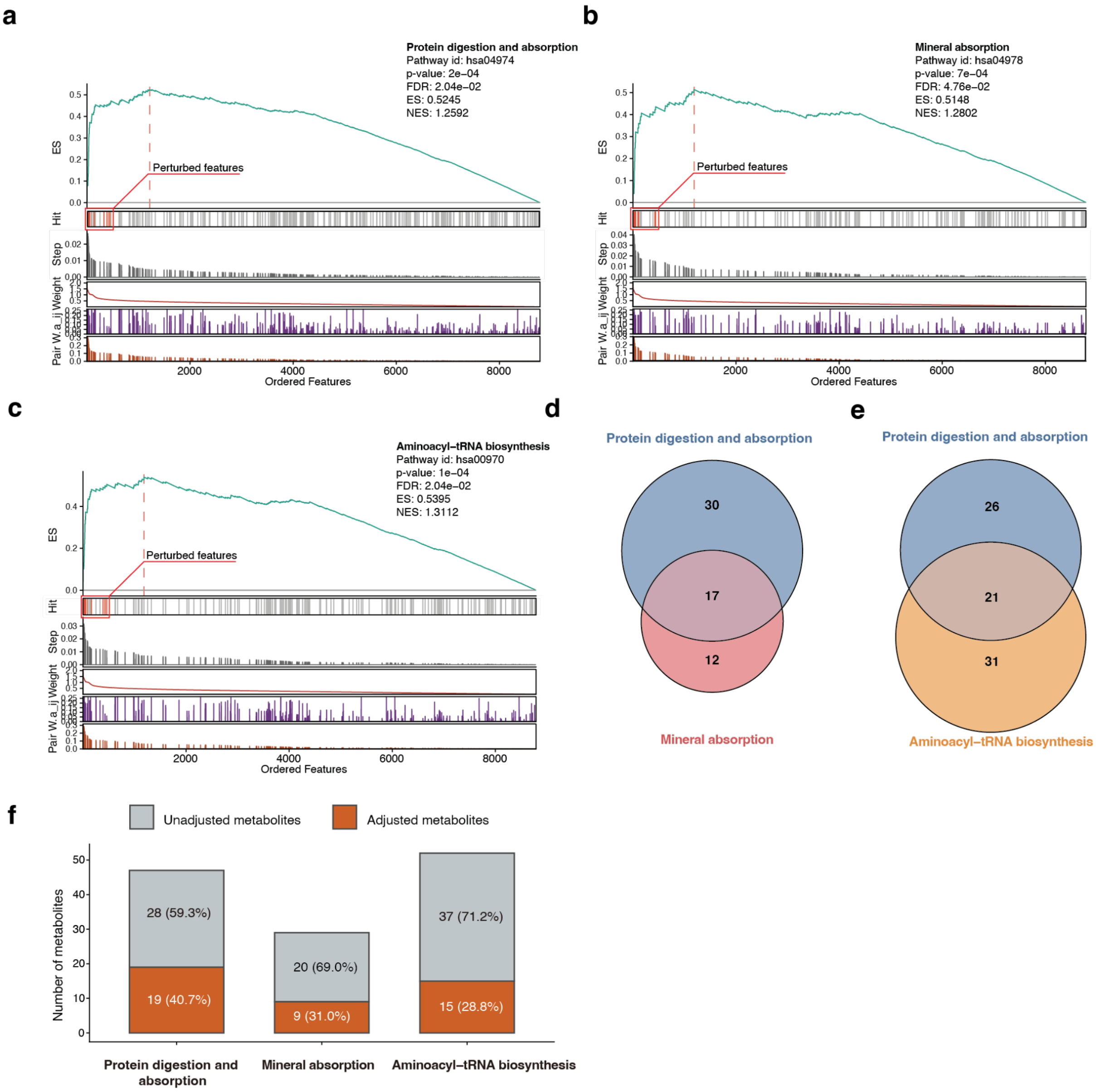


**Supplementary Figure 2 | Characterization of co-enriched metabolite sets in the protein digestion and absorption spike-in simulation. (a-c)** Enrichment plots for co-enriched metabolite sets in the spike-in simulations. **(d)** Overlap of metabolites between protein digestion and absorption and mineral absorption. **(e)** Overlap of metabolites between protein digestion and absorption and aminoacyl−tRNA biosynthesis. **(f)** Composition of adjusted and unadjusted metabolites within each enriched metabolite set.

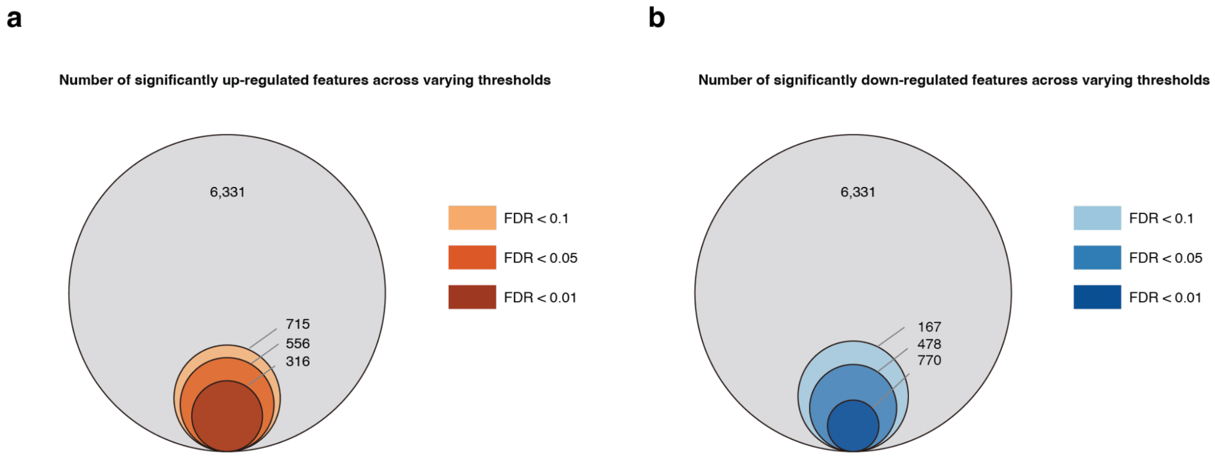


**Supplementary Figure 3 | Number of significant metabolic features across varying thresholds in the HD dataset. (a)** Number of significantly up-regulated features across varying thresholds. **(b)** Number of significantly down-regulated features across varying thresholds.

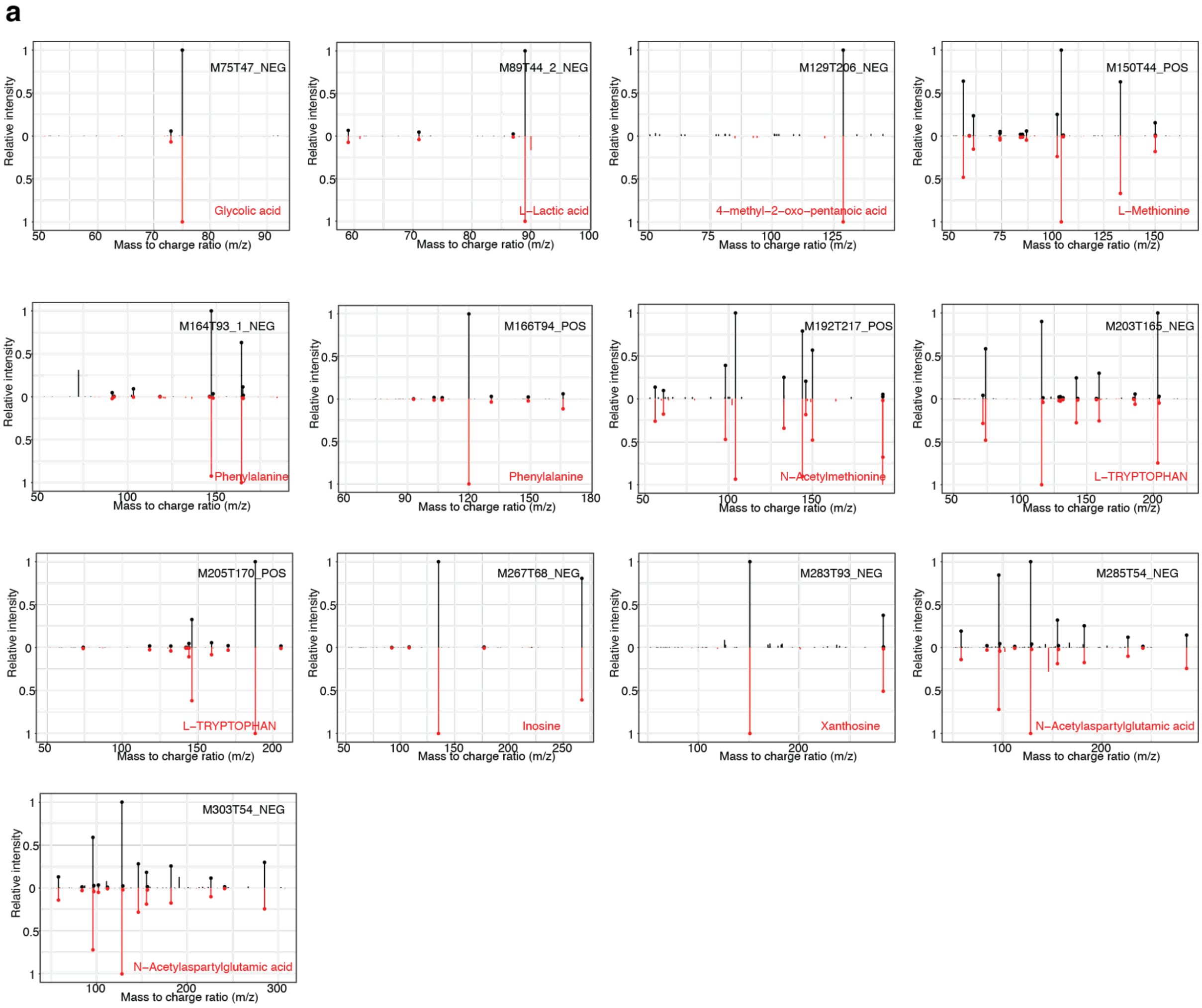


**Supplementary Figure 4 | Validation of Top 1 annotations by MS/MS spectra of standard compounds. (a)** MS/MS spectral matching between the top 1 annotations and the authentic standard compounds.

**a**

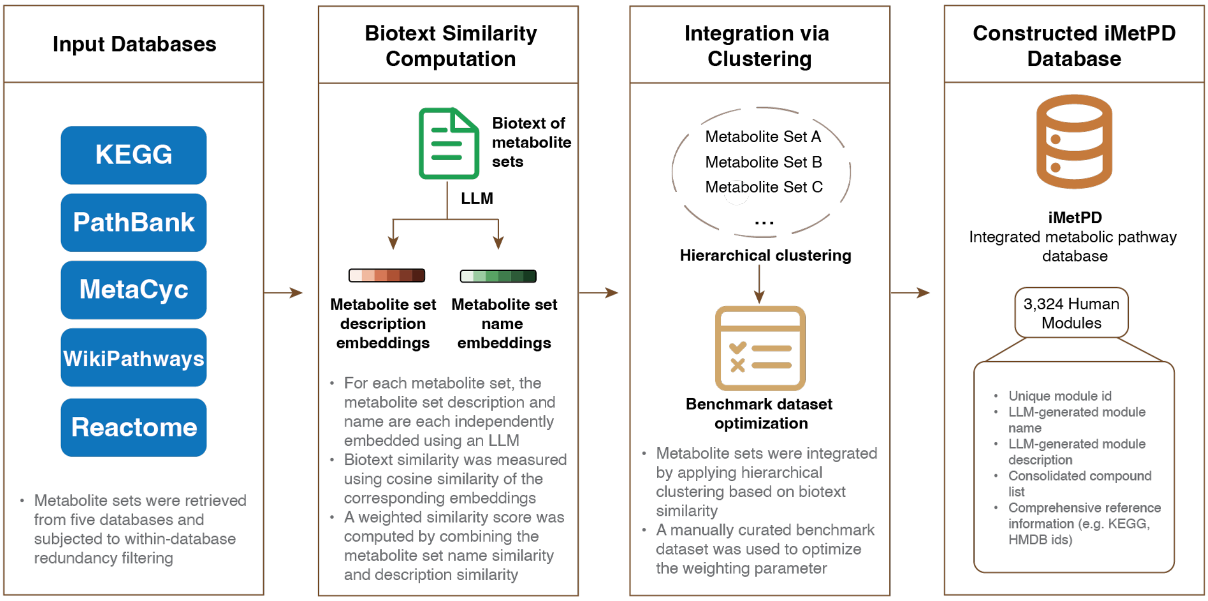


**Supplementary Figure 5 | Construction of the integrated metabolic pathway database (iMetPD) with five metabolic pathway databases. (a)** Construction of iMetPD based on metabolic pathways from KEGG, PathBank, MetaCyc, Wikipathways, and Reactome. Five metabolic pathway databases were downloaded, and redundant pathways within each database were removed. The remaining pathways were merged, and 4,913 metabolic pathways were obtained in total. Biotext embeddings for pathway description and pathway name were generated with LLM, and similarity between two metabolic pathways was determined based on pathway description similarity and pathway name similarity, optimized by a manually curated benchmark dataset. 3,324 integrated human metabolic modules were obtained through hierarchical clustering.

**Supplementary Tables**

**Supplementary Table 1. Confidence assessment of significantly enriched metabolite sets under correct matrix assignment**

| Pathway ID | Pathway name | ES | NES | p_value | FDR | Matrix confidence score | matrix_confidence_reason | Matrix source |
|---|---|---|---|---|---|---|---|---|
| hsa00100 | Steroid biosynthesis | 0.63227162 | 1.30537737 | 0.000999 | 0.01209901 | 75 | Plasma sterols and cholesterol precursors can reflect systemic cholesterol/steroid precursor metabolism and are commonly used as circulating biomarkers. Interpretation is still confounded by lipoprotein transport, diet, and tissue-specific synthesis. | plasma |
| hsa00140 | Steroid hormone biosynthesis | 0.74332423 | 1.52309079 | 0.000999 | 0.01209901 | 100 | Plasma is a highly appropriate matrix for steroid hormones because these compounds are secreted into circulation and routinely measured there. Levels can directly reflect endocrine steroid production, though timing and binding proteins affect interpretation. | plasma |
| hsa00524 | Neomycin, kanamycin and gentamicin biosynthesis | 0.60978531 | 1.25695419 | 0.000999 | 0.01209901 | 0 | Humans do not biosynthesize aminoglycoside antibiotics, so plasma metabolites would indicate exposure, medication, or contamination rather than pathway activity. This matrix is therefore unreliable for inferring this biosynthetic pathway. | plasma |
| hsa00830 | Retinol metabolism | 0.65985907 | 1.32376973 | 0.000999 | 0.01209901 | 50 | Plasma retinol and retinoid-related metabolites can provide some information on vitamin A transport and systemic status. However, homeostatic control, liver storage, diet, and binding proteins make pathway activity inference only moderate. | plasma |
| hsa00860 | Porphyrin metabolism | 0.64569703 | 1.2647581 | 0.000999 | 0.01209901 | 75 | Plasma porphyrins, bilirubin, and related heme catabolism products can reasonably reflect heme/porphyrin turnover and clinically relevant disturbances. Interpretation requires caution because liver clearance and hemolysis can strongly affect levels. | plasma |

| | | | | | | | | |
|---|---|---|---|---|---|---|---|---|
| hsa04913 | Ovarian steroidogenesis | 0.62774418 | 1.30296168 | 0.000999 | 0.01209901 | 75 | Plasma estradiol, progesterone, and androgens are useful circulating readouts of ovarian steroid output. Confidence is reduced by cycle phase, menopausal status, adrenal contribution, and exogenous hormones. | plasma |
| hsa04925 | Aldosterone synthesis and secretion | 0.77906991 | 1.55290961 | 0.000999 | 0.01209901 | 100 | Plasma aldosterone is a direct and standard clinical measure of adrenal mineralocorticoid secretion. Physiologic factors such as posture, sodium status, and renin activity influence interpretation but the matrix is highly appropriate. | plasma |
| hsa04976 | Bile secretion | 0.58512806 | 1.14757894 | 0.000999 | 0.01209901 | 75 | Plasma bile acids and bilirubin can indicate hepatic bile acid handling, cholestasis, and biliary transport. Levels also reflect enterohepatic cycling, meal timing, and liver clearance, so inference is good but not absolute. | plasma |
| hsa05200 | Pathways in cancer | 0.71839649 | 1.41745307 | 0.000999 | 0.01209901 | 25 | Plasma metabolites can show systemic cancer-associated metabolic changes, but this KEGG pathway is broad and dominated by signaling rather than a specific metabolic flux. Metabolite evidence is therefore weak and highly nonspecific. | plasma |

**Supplementary Table 2. Confidence assessment of significantly enriched metabolite sets under incorrect matrix assignment**

| Pathway ID | Pathway name | ES | NES | p_value | FDR | Matrix confidence score | matrix_confidence_reason | Matrix source |
|---|---|---|---|---|---|---|---|---|
| hsa00100 | Steroid biosynthesis | 0.63227162 | 1.30537737 | 0.000999 | 0.01209901 | 25 | Urine is not well suited for detecting nonpolar cholesterol/steroid biosynthetic intermediates, which are largely tissue- and lipoprotein-associated. Detected urinary steroids more often reflect downstream metabolism and excretion than core biosynthetic activity. | urine |
| hsa00140 | Steroid hormone biosynthesis | 0.74332423 | 1.52309079 | 0.000999 | 0.01209901 | 75 | Urinary steroid hormone metabolites are commonly used to infer systemic steroid production and endocrine activity. Interpretation still requires caution because renal handling, conjugation, and timing affect measured levels. | urine |
| hsa00524 | Neomycin, kanamycin and gentamicin biosynthesis | 0.60978531 | 1.25695419 | 0.000999 | 0.01209901 | 0 | This is a microbial antibiotic biosynthesis pathway, not a human urinary metabolic pathway. Detection in urine would most likely reflect drug exposure or excretion rather than biosynthetic activity. | urine |
| hsa00830 | Retinol metabolism | 0.65985907 | 1.32376973 | 0.000999 | 0.01209901 | 25 | Retinoid metabolism is mainly hepatic and tissue-associated, while urinary retinoids are limited and influenced by excretion or degradation. Urine therefore provides weak evidence for retinol pathway activity. | urine |
| hsa00860 | Porphyrin metabolism | 0.64569703 | 1.2647581 | 0.000999 | 0.01209901 | 75 | Urinary porphyrins and precursors such as ALA or PBG are clinically informative excretion markers of porphyrin/heme pathway disturbances. They can indicate altered pathway activity, though urine reflects accumulated excreted products rather than direct tissue flux. | urine |

| | | | | | | | | |
|---|---|---|---|---|---|---|---|---|
| hsa04913 | Ovarian steroidogenesis | 0.62774418 | 1.30296168 | 0.000999 | 0.01209901 | 50 | Urinary estrogen and progesterone metabolites can reflect ovarian steroid production, especially over integrated collection periods. Confidence is moderate because values are strongly affected by menstrual timing, sex, conjugation, and non-ovarian steroid sources. | urine |
| hsa04925 | Aldosterone synthesis and secretion | 0.77906991 | 1.55290961 | 0.000999 | 0.01209901 | 75 | Urinary aldosterone and its metabolites can reasonably reflect adrenal aldosterone production and are used in endocrine assessment. Interpretation is influenced by renal function, sodium balance, collection timing, and medication effects. | urine |
| hsa04976 | Bile secretion | 0.58512806 | 1.14757894 | 0.000999 | 0.01209901 | 50 | Urinary bile acids or bilirubin-related compounds may indicate altered hepatobiliary handling or cholestasis. However, urine reflects spillover and excretion rather than bile secretion itself, so pathway inference is indirect. | urine |
| hsa05200 | Pathways in cancer | 0.71839649 | 1.41745307 | 0.000999 | 0.01209901 | 25 | This pathway is broad and signaling-focused, while urinary metabolites provide nonspecific downstream or excretory changes. Urine alone gives weak confidence for inferring cancer pathway activity without supporting clinical or molecular data. | urine |

**Supplementary Table 3. 15 significant metabolite sets identified with FeatureMSEA before iterative refinement**

| pathway_id | pathway_name | KEGG_ID | ES | NES | p_value | FDR |
|---|---|---|---|---|---|---|
| hsa00020 | Citrate cycle (TCA cycle) | C00022{}C00024{}C00026{}C00036{}C00042{}C00068{}C00074{}C00091{}C00122{}C00149{}C00158{}C00311{}C00417{}C05125{}C05379{}C05381{}C15972{}C15973{}C16254{}C16255 | 0.6093 | 1.479 | 1.00E-04 | 0.00363 |
| hsa00040 | Pentose and glucuronate interconversions | C00022{}C00026{}C00029{}C00103{}C00111{}C00116{}C00167{}C00181{}C00191{}C00199{}C00204{}C00216{}C00231{}C00259{}C00266{}C00309{}C00310{}C00312{}C00333{}C00379{}C00433{}C00470{}C00474{}C00476{}C00502{}C00508{}C00514{}C00532{}C00558{}C00618{}C00714{}C00789{}C00800{}C00817{}C00905{}C01068{}C01101{}C01508{}C01904{}C02266{}C02273{}C02426{}C02753{}C03033{}C03291{}C03826{}C04053{}C04349{}C04575{}C05385{}C05411{}C05412{}C06118{}C06441{}C14899{}C15930{}C20680{}C22337 | 0.507 | 1.2038 | 0.0081 | 0.0803 |
| hsa00130 | Ubiquinone and other terpenoid-quinone biosynthesis | C00082{}C00156{}C00223{}C00251{}C00341{}C00353{}C00390{}C00423{}C00544{}C00811{}C00828{}C00885{}C01179{}C02059{}C02477{}C02483{}C02730{}C02949{}C03160{}C03313{}C03657{}C03964{}C03993{}C04145{}C05427{}C05807{}C05817{}C05819{}C05847{}C05848{}C05849{}C06986{}C10793{}C13309{}C14151{}C14152{}C14153{}C14154{}C14155{}C14156{}C15547{}C15882{}C15883{}C16519{}C16695{}C16999{}C17010{}C17017{}C17018{}C17412{}C17551{}C17552{}C17554{}C17559{}C17570{}C18131{}C18132{}C18133{}C18134{}C18135{}C19847{}C19858{}C19859{}C20265{}C20737{}C20738{}C20772{}C20773{}C21084{}C21860{}C22039 | 0.4769 | 1.2273 | 0.0053 | 0.0722 |
| hsa00250 | Alanine, aspartate and glutamate metabolism | C00014{}C00022{}C00025{}C00026{}C00036{}C00041{}C00042{}C00049{}C00064{}C00122{}C00152{}C00158{}C00169{}C00232{}C00334{}C00352{}C00402{}C00438{}C00940{}C01042{}C02362{}C03090{}C03406{}C03794{}C03912{}C12270{}C20775{}C20776 | 0.5666 | 1.4216 | 1.00E-04 | 0.00363 |
| hsa00280 | Valine, leucine and isoleucine degradation | C00024{}C00068{}C00091{}C00100{}C00123{}C00141{}C00164{}C00183{}C00233{}C00332{}C00349{}C00356{}C00407{}C00630{}C00671{}C00683{}C01205{}C01213{}C02170{}C02939{}C03069{}C03231{}C03284{}C03344{}C03345{}C03460{}C04405{}C05996{}C05998{}C06000{}C06001{}C06002{}C15972{}C15973{}C15974{}C15975{}C15976{}C15977{}C15978{}C15979{}C15980{}C20827 | 0.5547 | 1.2419 | 0.0095 | 0.0863 |

| | | | | | | |
|---|---|---|---|---|---|---|
| hsa00310 | Lysine degradation | C00024{}C00026{}C00037{}C00042{}C00047{}C00136{}C00164{}C00322{}C00332{}C00408{}C00431{}C00449{}C00450{}C00487{}C00489{}C00527{}C00739{}C00877{}C00956{}C00990{}C01142{}C01144{}C01149{}C01181{}C01186{}C01211{}C01259{}C01672{}C02188{}C02727{}C03087{}C03196{}C03239{}C03273{}C03366{}C03656{}C03793{}C04020{}C04076{}C04092{}C04487{}C05161{}C05231{}C05544{}C05545{}C05546{}C05548{}C05825{}C12455{}C16741 | 0.5014 | 1.1737 | 0.0119 | 0.0926 |
| hsa00350 | Tyrosine metabolism | C00022{}C00042{}C00082{}C00122{}C00146{}C00164{}C00232{}C00355{}C00483{}C00530{}C00544{}C00547{}C00628{}C00642{}C00788{}C00811{}C00822{}C01036{}C01060{}C01061{}C01161{}C01179{}C01384{}C01693{}C01829{}C01850{}C02167{}C02442{}C02465{}C02514{}C02515{}C03063{}C03077{}C03758{}C03765{}C03964{}C04043{}C04045{}C04052{}C04185{}C04186{}C04368{}C04642{}C04796{}C04797{}C05338{}C05350{}C05576{}C05577{}C05578{}C05579{}C05580{}C05581{}C05582{}C05583{}C05584{}C05585{}C05587{}C05588{}C05589{}C05593{}C05594{}C05595{}C05596{}C05600{}C05604{}C06044{}C06046{}C06047{}C06048{}C06199{}C06201{}C10447{}C17935{}C17936{}C17937{}C17938{}C22038 | 0.5082 | 1.3143 | 1.00E-04 | 0.00363 |
| hsa00360 | Phenylalanine metabolism | C00022{}C00024{}C00042{}C00079{}C00082{}C00084{}C00091{}C00122{}C00166{}C00180{}C00423{}C00512{}C00582{}C00596{}C00601{}C00642{}C00805{}C01198{}C01575{}C01586{}C01772{}C02137{}C02232{}C02265{}C02505{}C02763{}C02765{}C03519{}C03589{}C04044{}C04148{}C04351{}C04479{}C05332{}C05593{}C05598{}C05607{}C05629{}C05852{}C05853{}C06207{}C07086{}C08300{}C08301{}C11457{}C11588{}C12621{}C12622{}C12623{}C12624{}C14144{}C14145{}C15524{}C16719{}C17268{}C19945{}C19946{}C19975{}C20062 | 0.4621 | 1.1547 | 0.0132 | 0.0959 |
| hsa00400 | Phenylalanine, tyrosine and tryptophan biosynthesis | C00074{}C00078{}C00079{}C00082{}C00108{}C00119{}C00166{}C00251{}C00254{}C00279{}C00296{}C00354{}C00441{}C00463{}C00493{}C00587{}C00826{}C00944{}C01094{}C01179{}C01269{}C01302{}C02637{}C03175{}C03506{}C04302{}C04691{}C16848{}C16850{}C17235{}C20327{}C20653{}C20654{}C20710 | 0.4915 | 1.1982 | 0.0078 | 0.0803 |
| hsa00470 | D-Amino acid metabolism | C00022{}C00025{}C00026{}C00036{}C00037{}C00041{}C00047{}C00049{}C00062{}C00064{}C00065{}C00073{}C00077{}C00079{}C00084{}C00097{}C00133{}C00134{}C00135{}C00148{}C00166{}C00188{}C00217{}C00402{}C00431{}C00433{}C00515{}C00666{}C00680{}C00692{}C00739{}C00740{}C00763{}C00792{}C00793{}C00819{}C00820{}C00855{}C00993{}C01110{}C01157{}C01180{}C01667{}C01672{}C01726{}C02237{}C02265{}C02855{}C03239{}C03341{}C03440{}C03564{}C03771{}C0 | 0.4857 | 1.1601 | 0.0114 | 0.0926 |

| | | | | | | |
|---|---|---|---|---|---|---|
| | | 3933{}C03943{}C04260{}C04282{}C04457{}C05161{}C05620{}C05825{}C05939{}C05941{}C05942{}C06419{}C22024{}C22025 | | | | |
| hsa00630 | Glyoxylate and dicarboxylate metabolism | C00007{}C00011{}C00014{}C00022{}C00024{}C00025{}C00026{}C00027{}C00033{}C00036{}C00037{}C00042{}C00048{}C00058{}C00064{}C00065{}C00091{}C00100{}C00136{}C00149{}C00158{}C00160{}C00168{}C00197{}C00209{}C00258{}C00266{}C00311{}C00313{}C00332{}C00417{}C00552{}C00631{}C00683{}C00798{}C00877{}C00888{}C00898{}C00975{}C00988{}C01127{}C01146{}C01182{}C01213{}C01380{}C01732{}C01989{}C01990{}C02123{}C02405{}C03217{}C03459{}C03548{}C03561{}C03618{}C04348{}C06027{}C06028{}C06049{}C18026{}C18324{}C20238 | 0.5168 | 1.2181 | 0.0022 | 0.04 |
| hsa00740 | Riboflavin metabolism | C00016{}C00044{}C00061{}C00199{}C00235{}C00255{}C00474{}C01007{}C01268{}C01304{}C01352{}C01390{}C01727{}C01847{}C03114{}C04332{}C04454{}C04732{}C05995{}C15556{}C15563{}C18910{}C21214{}C21215 | 0.5642 | 1.4237 | 0.0013 | 0.0283 |
| hsa04742 | Taste transduction | C00002{}C00008{}C00020{}C00025{}C00031{}C00076{}C00080{}C00089{}C00095{}C00130{}C00133{}C00144{}C00149{}C00158{}C00208{}C00238{}C00334{}C00547{}C00575{}C00740{}C00780{}C01245{}C01327{}C01330{}C01451{}C01996{}C02265{}C06526{}C11045{}C12283{}C12284{}C19378{}D08836 | 0.5 | 1.2174 | 0.007 | 0.0803 |
| hsa04922 | Glucagon signaling pathway | C00022{}C00024{}C00026{}C00031{}C00036{}C00040{}C00042{}C00074{}C00076{}C00083{}C00085{}C00103{}C00122{}C00149{}C00158{}C00182{}C00186{}C00197{}C00311{}C00354{}C00575{}C00631{}C00665{}C00668{}C01172{}C01245 | 0.5157 | 1.2771 | 0.0044 | 0.0685 |
| hsa05230 | Central carbon metabolism in cancer | C00022{}C00024{}C00025{}C00026{}C00031{}C00036{}C00037{}C00041{}C00042{}C00049{}C00062{}C00064{}C00065{}C00073{}C00074{}C00078{}C00079{}C00082{}C00085{}C00092{}C00097{}C00122{}C00123{}C00135{}C00148{}C00149{}C00152{}C00158{}C00183{}C00186{}C00197{}C00311{}C00354{}C00407{}C00631{}C00665{}C00704 | 0.4934 | 1.2164 | 0.0012 | 0.0283 |

**Supplementary Table 4. 23 significant metabolite sets identified with FeatureMSEA after iterative refinement**

| pathway_id | pathway_name | KEGG_ID | ES | NES | p_value | FDR |
|---|---|---|---|---|---|---|
| hsa00020 | Citrate cycle (TCA cycle) | C00022{}C00024{}C00026{}C00036{}C00042{}C00068{}C00074{}C00091{}C00122{}C00149{}C00158{}C00311{}C00417{}C05125{}C05379{}C05381{}C15972{}C15973{}C16254{}C16255 | 0.5679 | 1.3566 | 0.0023 | 0.0139 |
| hsa00040 | Pentose and glucuronate interconversions | C00022{}C00026{}C00029{}C00103{}C00111{}C00116{}C00167{}C00181{}C00191{}C00199{}C00204{}C00216{}C00231{}C00259{}C00266{}C00309{}C00310{}C00312{}C00333{}C00379{}C00433{}C00470{}C00474{}C00476{}C00502{}C00508{}C00514{}C00532{}C00558{}C00618{}C00714{}C00789{}C00800{}C00817{}C00905{}C01068{}C01101{}C01508{}C01904{}C02266{}C02273{}C02426{}C02753{}C03033{}C03291{}C03826{}C04053{}C04349{}C04575{}C05385{}C05411{}C05412{}C06118{}C06441{}C14899{}C15930{}C20680{}C22337 | 0.5646 | 1.2793 | 1.00E-04 | 0.00182 |
| hsa00130 | Ubiquinone and other terpenoid-quinone biosynthesis | C00082{}C00156{}C00223{}C00251{}C00341{}C00353{}C00390{}C00423{}C00544{}C00811{}C00828{}C00885{}C01179{}C02059{}C02477{}C02483{}C02730{}C02949{}C03160{}C03313{}C03657{}C03964{}C03993{}C04145{}C05427{}C05807{}C05817{}C05819{}C05847{}C05848{}C05849{}C06986{}C10793{}C13309{}C14151{}C14152{}C14153{}C14154{}C14155{}C14156{}C15547{}C15882{}C15883{}C16519{}C16695{}C16999{}C17010{}C17017{}C17018{}C17412{}C17551{}C17552{}C17554{}C17559{}C17570{}C18131{}C18132{}C18133{}C18134{}C18135{}C19847{}C19858{}C19859{}C20265{}C20737{}C20738{}C20772{}C20773{}C21084{}C21860{}C22039 | 0.5505 | 1.2568 | 6.00E-04 | 0.00545 |
| hsa00190 | Oxidative phosphorylation | C00001{}C00002{}C00003{}C00004{}C00007{}C00008{}C00009{}C00013{}C00042{}C00061{}C00080{}C00122{}C00390{}C00399{}C00524{}C00536 | 0.6544 | 1.4825 | 0.004 | 0.0208 |
| hsa00220 | Arginine biosynthesis | C00011{}C00014{}C00025{}C00026{}C00049{}C00062{}C00064{}C00077{}C00086{}C00122{}C00169{}C00327{}C00437{}C00624{}C01010{}C01250{}C03406{}C04133{}C15532{}C20948{}C20949{}C20950{}C20951 | 0.5959 | 1.3086 | 4.00E-04 | 0.00484 |
| hsa00250 | Alanine, aspartate and glutamate metabolism | C00014{}C00022{}C00025{}C00026{}C00036{}C00041{}C00042{}C00049{}C00064{}C00122{}C00152{}C00158{}C00169{}C00232{}C00334{}C00352{}C00402{}C00438{}C00940{}C01042{}C02362{}C03090{}C03406{}C03794{}C03912{}C12270{}C20775{}C20776 | 0.6037 | 1.4558 | 1.00E-04 | 0.00182 |

| | | | | | | |
|---|---|---|---|---|---|---|
| hsa00260 | Glycine, serine and threonine metabolism | C00011{}C00014{}C00022{}C00037{}C00048{}C00049{}C00065{}C00078{}C00097{}C00101{}C00109{}C00114{}C00143{}C00168{}C00188{}C00197{}C00213{}C00258{}C00263{}C00300{}C00430{}C00441{}C00546{}C00576{}C00581{}C00631{}C00719{}C00740{}C00986{}C01005{}C01026{}C01102{}C01242{}C01888{}C02051{}C02291{}C02737{}C02972{}C03082{}C03194{}C03232{}C03283{}C03508{}C05519{}C06231{}C06442{}C16432{}C19929 | 0.5473 | 1.1998 | 8.00E-04 | 0.00545 |
| hsa00280 | Valine, leucine and isoleucine degradation | C00024{}C00068{}C00091{}C00100{}C00123{}C00141{}C00164{}C00183{}C00233{}C00332{}C00349{}C00356{}C00407{}C00630{}C00671{}C00683{}C01205{}C01213{}C02170{}C02939{}C03069{}C03231{}C03284{}C03344{}C03345{}C03460{}C04405{}C05996{}C05998{}C06000{}C06001{}C06002{}C15972{}C15973{}C15974{}C15975{}C15976{}C15977{}C15978{}C15979{}C15980{}C20827 | 0.6207 | 1.2781 | 6.00E-04 | 0.00545 |
| hsa00310 | Lysine degradation | C00024{}C00026{}C00037{}C00042{}C00047{}C00136{}C00164{}C00322{}C00332{}C00408{}C00431{}C00449{}C00450{}C00487{}C00489{}C00527{}C00739{}C00877{}C00956{}C00990{}C01142{}C01144{}C01149{}C01181{}C01186{}C01211{}C01259{}C01672{}C02188{}C02727{}C03087{}C03196{}C03239{}C03273{}C03366{}C03656{}C03793{}C04020{}C04076{}C04092{}C04487{}C05161{}C05231{}C05544{}C05545{}C05546{}C05548{}C05825{}C12455{}C16741 | 0.5764 | 1.219 | 7.00E-04 | 0.00545 |
| hsa00330 | Arginine and proline metabolism | C00012{}C00019{}C00022{}C00025{}C00048{}C00062{}C00077{}C00086{}C00134{}C00148{}C00179{}C00213{}C00223{}C00300{}C00315{}C00334{}C00406{}C00431{}C00436{}C00441{}C00533{}C00555{}C00581{}C00750{}C00763{}C00791{}C00884{}C00986{}C01035{}C01043{}C01110{}C01137{}C01157{}C01165{}C01250{}C01682{}C01877{}C02305{}C02565{}C02647{}C02714{}C02946{}C03078{}C03166{}C03287{}C03296{}C03375{}C03415{}C03564{}C03771{}C03912{}C04137{}C04281{}C04498{}C05147{}C05931{}C05932{}C05933{}C05936{}C05938{}C05945{}C05946{}C05947{}C10497{}C15699{}C15700{}C15767{}C18172{}C18174{}C18325{}C18326{}C19706 | 0.5426 | 1.2082 | 1.00E-04 | 0.00182 |
| hsa00350 | Tyrosine metabolism | C00022{}C00042{}C00082{}C00122{}C00146{}C00164{}C00232{}C00355{}C00483{}C00530{}C00544{}C00547{}C00628{}C00642{}C00788{}C00811{}C00822{}C01036{}C01060{}C01061{}C01161{}C01179{}C01384{}C01693{}C01829{}C01850{}C02167{}C02442{}C02465{}C02514{}C02515{}C03063{}C03077{}C03758{}C03765{}C03964{}C04043{}C04045{}C04052{}C04185{}C04186{}C04368{}C04642{}C04796{}C04797{}C05338{}C05350{}C05576{}C05577{}C05578{}C05579{}C05580{}C05581{}C05582{}C05583{}C05584{}C05585{}C05587{}C05588{}C05589{}C05593{}C05594{}C05595{}C05596{}C05600{}C05604{}C06044{}C06046{}C06047{}C06048{}C06199{}C06201{}C10447{}C17935{}C17936{}C17937{}C17938{}C22038 | 0.5929 | 1.351 | 1.00E-04 | 0.00182 |

| | | | | | | |
|---|---|---|---|---|---|---|
| hsa00360 | Phenylalanine metabolism | C00022{}C00024{}C00042{}C00079{}C00082{}C00084{}C00091{}C00122{}C00166{}C00180{}C00423{}C00512{}C00582{}C00596{}C00601{}C00642{}C00805{}C01198{}C01575{}C01586{}C01772{}C02137{}C02232{}C02265{}C02505{}C02763{}C02765{}C03519{}C03589{}C04044{}C04148{}C04351{}C04479{}C05332{}C05593{}C05598{}C05607{}C05629{}C05852{}C05853{}C06207{}C07086{}C08300{}C08301{}C11457{}C11588{}C12621{}C12622{}C12623{}C12624{}C14144{}C14145{}C15524{}C16719{}C17268{}C19945{}C19946{}C19975{}C20062 | 0.5405 | 1.2175 | 6.00E-04 | 0.00545 |
| hsa00400 | Phenylalanine, tyrosine and tryptophan biosynthesis | C00074{}C00078{}C00079{}C00082{}C00108{}C00119{}C00166{}C00251{}C00254{}C00279{}C00296{}C00354{}C00441{}C00463{}C00493{}C00587{}C00826{}C00944{}C01094{}C01179{}C01269{}C01302{}C02637{}C03175{}C03506{}C04302{}C04691{}C16848{}C16850{}C17235{}C20327{}C20653{}C20654{}C20710 | 0.5494 | 1.2344 | 8.00E-04 | 0.00545 |
| hsa00470 | D-Amino acid metabolism | C00022{}C00025{}C00026{}C00036{}C00037{}C00041{}C00047{}C00049{}C00062{}C00064{}C00065{}C00073{}C00077{}C00079{}C00084{}C00097{}C00133{}C00134{}C00135{}C00148{}C00166{}C00188{}C00217{}C00402{}C00431{}C00433{}C00515{}C00666{}C00680{}C00692{}C00739{}C00740{}C00763{}C00792{}C00793{}C00819{}C00820{}C00855{}C00993{}C01110{}C01157{}C01180{}C01667{}C01672{}C01726{}C02237{}C02265{}C02855{}C03239{}C03341{}C03440{}C03564{}C03771{}C03933{}C03943{}C04260{}C04282{}C04457{}C05161{}C05620{}C05825{}C05939{}C05941{}C05942{}C06419{}C22024{}C22025 | 0.5426 | 1.2016 | 7.00E-04 | 0.00545 |
| hsa00630 | Glyoxylate and dicarboxylate metabolism | C00007{}C00011{}C00014{}C00022{}C00024{}C00025{}C00026{}C00027{}C00033{}C00036{}C00037{}C00042{}C00048{}C00058{}C00064{}C00065{}C00091{}C00100{}C00136{}C00149{}C00158{}C00160{}C00168{}C00197{}C00209{}C00258{}C00266{}C00311{}C00313{}C00332{}C00417{}C00552{}C00631{}C00683{}C00798{}C00877{}C00888{}C00898{}C00975{}C00988{}C01127{}C01146{}C01182{}C01213{}C01380{}C01732{}C01989{}C01990{}C02123{}C02405{}C03217{}C03459{}C03548{}C03561{}C03618{}C04348{}C06027{}C06028{}C06049{}C18026{}C18324{}C20238 | 0.5762 | 1.2959 | 1.00E-04 | 0.00182 |
| hsa00740 | Riboflavin metabolism | C00016{}C00044{}C00061{}C00199{}C00235{}C00255{}C00474{}C01007{}C01268{}C01304{}C01352{}C01390{}C01727{}C01847{}C03114{}C04332{}C04454{}C04732{}C05995{}C15556{}C15563{}C18910{}C21214{}C21215 | 0.7191 | 1.4658 | 2.00E-04 | 0.00311 |
| hsa00860 | Porphyrin metabolism | C00025{}C00032{}C00037{}C00188{}C00194{}C00430{}C00486{}C00500{}C00524{}C00541{}C00748{}C00853{}C00931{}C00992{}C01024{}C01051{}C01079{}C01708{}C02139{}C02191{}C02463{}C02469{}C02800{}C02823{}C02880{}C02987{}C03029{}C03114{}C03179{}C03194{}C03263{}C03373{}C03516{}C03741{}C04122{}C04536{}C04778{}C05306{}C05307{} | 0.5723 | 1.2398 | 0.007 | 0.0347 |

| | | | | | | |
|---|---|---|---|---|---|---|
| | | C05766{}C05767{}C05768{}C05769{}C05770{}C05772{}C05773{}C05774{}C05775{}C05777{}C05778{}C05779{}C05780{}C05781{}C05782{}C05783{}C05784{}C05785{}C05786{}C05787{}C05789{}C05790{}C05791{}C05793{}C05794{}C05795{}C05797{}C05798{}C05912{}C05913{}C06319{}C06320{}C06399{}C06406{}C06407{}C06408{}C06416{}C06503{}C06504{}C06505{}C06506{}C06507{}C06508{}C06509{}C06510{}C11242{}C11243{}C11538{}C11540{}C11542{}C11543{}C11545{}C11630{}C11829{}C11830{}C11831{}C11832{}C11850{}C11851{}C12147{}C14818{}C14819{}C15670{}C15672{}C16242{}C16243{}C16244{}C16540{}C16541{}C17401{}C18021{}C18022{}C18064{}C18098{}C18151{}C18152{}C18153{}C18154{}C18155{}C18156{}C18157{}C18160{}C18161{}C18162{}C18163{}C19608{}C20666{}C21217{}C21284{}C21427{}C21428{}C21429{}C21431{}C21432{}C21433{}C21434{}C21435{}C21510{}C21511{}C21512{}C21582{}C21723{}C21724{}C21764{}C21835{}C22338{}C22339{}C22450{}C22451 | | | | |
| hsa02010 | ABC transporters | C00009{}C00025{}C00031{}C00032{}C00034{}C00037{}C00038{}C00041{}C00047{}C00049{}C00051{}C00059{}C00062{}C00064{}C00065{}C00070{}C00077{}C00079{}C00086{}C00088{}C00089{}C00093{}C00095{}C00098{}C00107{}C00114{}C00116{}C00120{}C00121{}C00123{}C00134{}C00135{}C00137{}C00140{}C00148{}C00151{}C00159{}C00175{}C00181{}C00183{}C00185{}C00188{}C00208{}C00212{}C00243{}C00244{}C00245{}C00255{}C00259{}C00288{}C00291{}C00294{}C00299{}C00315{}C00320{}C00330{}C00333{}C00338{}C00378{}C00379{}C00387{}C00392{}C00407{}C00430{}C00470{}C00475{}C00487{}C00491{}C00492{}C00503{}C00526{}C00559{}C00719{}C00794{}C00855{}C00865{}C00881{}C00919{}C00973{}C01083{}C01153{}C01157{}C01177{}C01181{}C01279{}C01330{}C01417{}C01487{}C01606{}C01630{}C01667{}C01674{}C01682{}C01684{}C01762{}C01834{}C01835{}C01935{}C01946{}C02160{}C02273{}C03557{}C03611{}C03619{}C04114{}C04137{}C05349{}C05402{}C05512{}C05776{}C06227{}C06229{}C06230{}C06232{}C06687{}C06705{}C06706{}C06707{}C06767{}C07662{}C07663{}C11612{}C13768{}C14818{}C14819{}C15521{}C15719{}C16421{}C16692{}C19609{}C19872{}C20570{}C20571{}C20572{}C20573{}C20679{}C21066{}C22040{}G00457 | 0.4822 | 1.1112 | 0.0174 | 0.0825 |
| hsa04024 | cAMP signaling pathway | C00020{}C00042{}C00076{}C00080{}C00165{}C00186{}C00212{}C00238{}C00288{}C00334{}C00416{}C00547{}C00575{}C00584{}C00698{}C00780{}C00788{}C01089{}C01245{}C01312{}C01330{}C01996{}C03758{}C20792{}C20793 | 0.5529 | 1.252 | 0.0025 | 0.0143 |

| | | | | | | |
|---|---|---|---|---|---|---|
| hsa04080 | Neuroactive ligand-receptor interaction | C00002{}C00008{}C00015{}C00025{}C00037{}C00049{}C00075{}C00099{}C00212{}C00245{}C00334{}C00388{}C00398{}C00483{}C00506{}C00547{}C00584{}C00639{}C00681{}C00696{}C00735{}C00780{}C00788{}C01120{}C01312{}C01516{}C01598{}C01829{}C01996{}C02165{}C02166{}C02198{}C02465{}C03758{}C04227{}C04598{}C05113{}C05951{}C05952{}C06124{}C06314{}C06865{}C11695{}C12270{}C12271{}C12272{}C13856{}C15890{}C15891{}C15995{}C16511{}C16512 | 0.5724 | 1.2686 | 1.00E-04 | 0.00182 |
| hsa04742 | Taste transduction | C00002{}C00008{}C00020{}C00025{}C00031{}C00076{}C00080{}C00089{}C00095{}C00130{}C00133{}C00144{}C00149{}C00158{}C00208{}C00238{}C00334{}C00547{}C00575{}C00740{}C00780{}C01245{}C01327{}C01330{}C01451{}C01996{}C02265{}C06526{}C11045{}C12283{}C12284{}C19378{}D08836 | 0.5455 | 1.2714 | 0.001 | 0.00641 |
| hsa04922 | Glucagon signaling pathway | C00022{}C00024{}C00026{}C00031{}C00036{}C00040{}C00042{}C00074{}C00076{}C00083{}C00085{}C00103{}C00122{}C00149{}C00158{}C00182{}C00186{}C00197{}C00311{}C00354{}C00575{}C00631{}C00665{}C00668{}C01172{}C01245 | 0.5338 | 1.2856 | 0.0029 | 0.0158 |
| hsa05230 | Central carbon metabolism in cancer | C00022{}C00024{}C00025{}C00026{}C00031{}C00036{}C00037{}C00041{}C00042{}C00049{}C00062{}C00064{}C00065{}C00073{}C00074{}C00078{}C00079{}C00082{}C00085{}C00092{}C00097{}C00122{}C00123{}C00135{}C00148{}C00149{}C00152{}C00158{}C00183{}C00186{}C00197{}C00311{}C00354{}C00407{}C00631{}C00665{}C00704 | 0.5239 | 1.239 | 4.00E-04 | 0.00484 |

**Supplementary Table 5. Scoring criteria for metabolite class annotation confidence**

| Item | Rule | Score | Reason |
|---|---|---|---|
| MS2 annotation | Any feature's annotation is based on the MS2 spectrum | 150 | MS/MS spectral confirmation is the strongest evidence for compound identity (similar to Level 2 or 1 in the Metabolomics Standards Initiative). It deserves the highest score. |
| Adduct types | [M+H]+ | 50 | Protonated or deprotonated ions are the most common and most reliable signals in LC-MS. Common adducts (like Na+, NH4+) provide supportive evidence but are less reliable than the main molecular ion. Rare adducts get lower scores. |
| | [M+Na]+ | 30 | |
| | [M+NH4]+ | 30 | |
| | [M+K]+ | 20 | |
| | Other positive mode ion type | 10 | |
| | [M−H]− | 50 | |
| | [M+Cl]− | 30 | |
| | [M+FA]− | 30 | |
| | Other negative mode ion type | 10 | |
| Isotope Peaks | Each detected M+1 peak | 10 | The presence of M+1 peaks confirms isotopic patterns (mainly due to natural 13C). Observing this pattern is good evidence that the peak is real and not noise. |

| | | | |
|---|---|---|---|
| | Each detected M+2 peak; +5 points only if the corresponding M+1 peak is also present.<br>If M+2 is present but M+1 is absent, do not add any score for M+2. In fact, such a scenario is suspicious and may indicate a noise peak or misannotation, because M+2 is rarely visible without M+1. | 5 | M+2 peaks often arise from multiple 13C atoms or from heavier isotopes (e.g. ^37Cl). It's extremely rare to see an M+2 peak in the absence of M+1. Therefore, we add points for M+2 only when isotopic patterns appear complete. |
| Co-elution and peak correlation | Correlated intensity profiles or correlated peak shapes | NA | If features co-elute closely and show similar intensity patterns across samples, they are likely from the same compound (e.g. different adducts, isotopes, fragments). This is a principle used in tools like CAMERA and GNPS feature networking. |

**Supplementary Note**

**Multi-evidence feature-to-metabolite annotation**

Features are first annotated with the MS1 database (KEGG[1] or HMDB[2]) using accurate mass matching (15 ppm), considering common adduct types in the corresponding ion mode. Features are then grouped into metabolite feature clusters (MFCs) based on chemical identity and retention time, with a tolerance of 10s by default. Within each MFC, isotope peaks ([M+1], [M+2], [M+3]) are identified based on predicted theoretical isotope distributions, with a mass tolerance of 15 ppm and a retention time tolerance of 10s.

A confidence score is assigned to each MFC by summing evidence-based scores across its member features according to predefined criteria (**Supplementary Table 5**). Briefly, MS/MS-based annotations contribute 150 points; primary adducts ($[M+H]^+$ or $[M-H]^-$) contribute 50 points; secondary adducts contribute 10–30 points; each detected isotope peak contributes 10 points, with [M+2] scored only when the corresponding [M+1] is also present. The theoretical maximum score is approximately 350 points for a metabolite detected in both ion modes with MS/MS confirmation.

To resolve annotation conflicts and remove redundancy, MFCs are pruned iteratively. If a metabolite maps to multiple MFCs, those scoring less than half of the highest-scoring MFC for the metabolite are removed. If a feature belongs to multiple MFCs with different scores, the annotations to the lower-scoring MFC are removed. This pruning is repeated until convergence. The final annotation table includes all remaining candidates with their confidence scores and serves as the feature annotation score table for feature-level enrichment scoring.

**LLM-assisted biological plausibility assessment**

*Matrix confidence assessment.* The system prompt consisted of three components:

(1) Task description: "You're an expert in LC-MS untargeted metabolomics pathway interpretation. Task: For each pathway, assess the CONFIDENCE of inferring pathway activity from metabolites detected in the given Sample Source (matrix). Note that pathways do not exist directly in biological matrices, but their activity can be inferred from the presence and levels of related metabolites."

(2) Scoring criteria: "Use DISCRETE ANCHORS: matrix_confidence (0/25/50/75/100): 100: High confidence - metabolites in this matrix reliably indicate pathway activity. 75: Good confidence - metabolites likely reflect pathway activity with reasonable certainty. 50: Moderate confidence - metabolites may indicate pathway activity, but interpretation requires caution. 25: Low confidence - metabolites provide weak evidence of pathway activity due to confounding factors. 0: No confidence - metabolites in this matrix are unreliable indicators of pathway activity."

(3) Matrix-specific hard constraints: "Matrix interpretation guidelines (STRICT CONSTRAINTS): Urine: Reflects excretion and filtration products; good for catabolic end-products. STRICT CONSTRAINT: Score matrix_confidence strictly as 0 for 'Aminoacyl-tRNA biosynthesis' and 'Amino acid biosynthesis'. Amino acids in urine typically result from filtration/pathological leaks rather than reflecting active biosynthesis, making inference unreliable. Feces: Dominated by microbial metabolism and dietary remnants. STRICT CONSTRAINT: Score matrix_confidence strictly as 0 for 'Lipid biosynthesis' and host 'Amino acid metabolism'. Fecal metabolites are primarily microbial or dietary in origin, not reliable indicators of host metabolic pathway activity. Plasma/Serum/Blood: Reflects systemic circulation and active metabolism. STRICT CONSTRAINT: Score matrix_confidence strictly as 0 for 'Amino acid

biosynthesis'. Blood amino acids reflect consumption/turnover rather than active biosynthesis, making inference of biosynthetic pathway activity unreliable."

*Topic relevance assessment.* For metabolite sets without any literature support from PubMed, the following LLM fallback prompt was used, consisting of two components:

(1) Task description: "You are an expert in metabolomics, biochemistry, and molecular biology. Task: For each metabolic pathway listed below, score how likely this pathway is to be relevant to the research topic."

(2) Scoring criteria: "Use DISCRETE ANCHORS for topic_confidence_score (0/25/50/75/100): 100: Strong, well-established link between this pathway and the research topic. 75: Likely relevant; credible biochemical or clinical connections exist. 50: Possibly relevant; indirect or context-dependent connections. 25: Unlikely relevant; only weak or speculative connections. 0: No meaningful connection to the research topic."

(3) Evaluation dimensions: "Consider: known biochemical links between the pathway and the disease/condition; potential upstream/downstream regulatory connections; shared metabolites or cofactors; published hypotheses or emerging evidence."

**Construction of the integrated metabolic pathway database (iMetPD)**

Metabolic set entries were retrieved from five databases: KEGG[1], PathBank[3], MetaCyc[4], WikiPathways[5], and Reactome[6] (**Supplementary Fig. 5**). For each database, redundant metabolite sets with identical descriptions were removed prior to integration. Essential metabolite set attributes were extracted, including metabolite set identifiers, names, descriptions, compound lists, and classification information. Metabolite identifiers across databases were converted to KEGG and HMDB formats using the MetOrigin database in TidyMass2[7]. A strict correspondence was kept between metabolite names and identifiers by preserving the order of entries within the compound list of each metabolite set.

Biotext similarity between two metabolite sets was defined based on the cosine similarity of embeddings generated from metabolite set names and descriptions using text-embedding-3-small. A weighted similarity score based on metabolite set name similarity and description similarity was computed for hierarchical clustering. The optimal weighting parameter was determined with a manually curated benchmark dataset. Metabolite sets assigned to the same cluster were merged into a single module, with compound lists consolidated across all member metabolite sets. GPT-4o was used to generate a unified name and description for each module based on the names and descriptions of its constituent metabolite sets (**Supplementary Fig. 5**).

The resulting iMetPD contains 3,324 human metabolic modules, including detailed information such as unique module identifier, LLM-generated module name and description, consolidated compound list, and comprehensive cross-database reference information including KEGG and HMDB identifiers.
